# Species- and strain-effects in interferon, innate immunity, and barrier transcriptional response of microbiome interactions of 3D respiratory epithelial cultures

**DOI:** 10.1101/2023.09.25.559418

**Authors:** Mian Horvath, Ruoyu Yang, Diana Cadena Castaneda, Megan Callender, Elizabeth Fleming, Anita Voigt, Ryan Caldwell, José Fachi, Blanda Di Luccia, Zoe Scholar, Peter Yu, Andrew Salner, Marco Colonna, Karolina Palucka, Julia Oh

**Affiliations:** The Jackson Laboratory for Genomic Medicine, Farmington, CT, 06032, USA; UCONN Health (University of Connecticut), Farmington, CT, 06030, USA; Washington University School of Medicine, St. Louis, MO, 63110, USA; Hartford Healthcare Cancer Institute, Hartford, CT, 06102, USA

## Abstract

The microbiome modulates the respiratory epithelium’s immunomodulatory functions. To explore how the microbiome’s biodiversity affects microbe-epithelial interactions, we screened 58 phylogenetically diverse microbes for their transcriptomic effect on tracheobronchial air-liquid interface cell (ALI) cultures. We found distinct species- and strain-level differences in host innate immunity, and epithelial barrier response. Strikingly, we found that host interferon, an antiviral immune response, was one of the most variable host processes, and this variability was not driven by microbial phylogenetic diversity, bioburden, nor by the microbe’s ability to stimulate other innate immunity pathways. Our study provides a foundation for understanding how the respiratory microbiome’s biodiversity affects epithelial, and particularly, antiviral innate immunity.

## Introduction

The respiratory epithelium is the first line of defense against potentially damaging environmental stimuli, such as allergens, toxins, and microbial pathogens. In addition to being a highly specialized physical barrier, regulating what can enter systemic circulation and removing foreign particles, it can directly contribute to innate immunity^1,2^. For example, the respiratory epithelium produces antimicrobial compounds (e.g., lysozymes^3^, defensins^4,5^, and cathelicidins^6^) and numerous cytokines, including interferons which have direct antiviral functions^7^ and other cytokines that recruit immune cells^8–10^. Understanding factors that modulate homeostasis and respiratory epithelial response to diverse stimuli is critical to developing new therapies that address infectious or chronic respiratory diseases.

A major modulator of the epithelium and its immunomodulatory functions is the microbiome, the diverse bacteria, fungi, and viruses that colonize the human body. Microbial colonization in the respiratory tract varies significantly between different regions of the respiratory tract due to differing environmental conditions. For example, the nares more closely resemble the skin in terms of species’ prevalence, with *Staphylococcus* (*S*.), *Corynebacterium*, and *Dolosigranulum* as some of the predominant genera. The microbes colonizing the oropharynx resemble oral microbes, with genera like *Streptococcus* (*Str*.), *Neisseria*, *Prevotella*, and *Rothia*. Finally, the lung microbiome is predominantly anaerobic genera such as *Prevotella, Str.*, and *Veillonella*^11^.

While microbial colonization is known to modulate the respiratory epithelium and subsequent immune responses, the effects of microbial colonization can be species- and strain-dependent, such that different microbial isolates of the same genus or species can have divergent effects on the host. For example, colonization with patient-derived *S. epidermidis*, but not with a lab strain, decreased influenza A infection^12^. Another study showed that colonization with *Str*. *salivarius*, but not *S*. *epidermidis*, increased secretion of numerous cytokines in an allergic rhinitis model^13^. While few studies have investigated the effect of microbial colonization on respiratory epithelial barrier integrity, in colonic epithelium, *Lactobacillus* (*L*.) *rhamnosus*, but not *L*. *crispatus*, prevented epithelial barrier dysfunction^14^.

However, our understanding of the mechanisms by which the respiratory microbiome modulates epithelial immunity, and potentially barrier functions, is still in its infancy compared to our understanding of gut microbiome-epithelial interactions. The above studies and others^15–18^ have only examined a handful of microbial isolates, which offers a limited recapitulation of the biodiversity seen in the respiratory tract and limits our understanding of the preventative or synergistic role of the microbiome in infection response as well as potential development of microbiome-based therapies that could enhance immune response or barrier integrity. One barrier to systematically investigating microbe-epithelial interactions is that few models can accommodate microbial diversity. For example, murine models, while possessing a full repertoire of innate and adaptive immune elements, are relatively low-throughput and possess endogenous microbiota that can differ significantly from species in human microbiomes^19^. While enabling throughput, 2D *in vitro* human cell culture screens with epithelial monolayers cannot replicate the cellular complexity as well as the inherently spatially constrained interactions with the microbiome that is observed in the native respiratory epithelium.

Here, we leverage 3D air-liquid interface cultures (ALI) to address spatial aspects of microbiome-epithelial interactions with sufficient throughput to interrogate a range of diverse microbiota. ALIs are a flat, multi-cell type, multi-layered epithelium, derived from epithelial progenitor cells, with the apical surface exposed to the air. They are morphologically^20,21^ and transcriptionally^22^ more similar to *in vivo* epithelium than 2D cultures, and, importantly, provide the necessary throughput to examine the effects of colonization of a diversity of microbiota. We colonized ALIs with 58 phylogenetically diverse, patient-derived respiratory microbes to examine the effect of microbial colonization on the transcriptional profile of tracheobronchial epithelium. Global host transcriptional response was measured with RNA-seq at two timepoints following colonization, with the overarching goal of reconstructing how activation of host processes, particularly antiviral innate immunity, differed based on microbial phylogeny. We identified notable species- and strain-level differences in host innate immunity and epithelial barrier integrity responses. Strikingly, we found the most variability in host interferon induction, a canonically antiviral immune response. Furthermore, a microbe’s propensity to induce interferon was not necessarily linked to its potential to activate antibacterial innate immunity. Taken together, our study provides a rich dataset on how microbial phylogenetic diversity affects microbiome modulation of host antiviral innate immunity, antibacterial innate immunity, and barrier responses in a 3D respiratory epithelial model.

## Results

### Microbiome Cultivation

As part of our pipeline to cultivate respiratory microbiota for screening, we collected samples from 47 patients (ages ranging from 46-83 years) who were recruited as part of a broader study investigating immunological changes in lung cancer. The patients were sampled at the tongue dorsum and nares as non-invasive approximations of the respiratory tract^23,24^ (Fig. 1A). While we emphasize that our goal in this study was not to categorically identify differences in this lung cancer cohort, we characterized the metagenome of these patient samples. In the nares, a few genera (i.e., *Cutibacterium*, *Corynebacterium*, and *Staphylococcus*) comprised the majority of identified microbes, which was concordant with other characterizations of healthy adult nasal microbiomes^25,26^ (Fig. 1B). Tongue dorsum samples were more diverse, including *Rothia mucilaginosa*, *Prevotella* spp., *Streptococcus* spp., *Veillonella* spp., *Neisseria subflava*, and *Haemophilus parainfluenzae*, which have previously been reported as highly abundant at this oral site^25,27^ (Fig. 1C).

**Figure 1:**
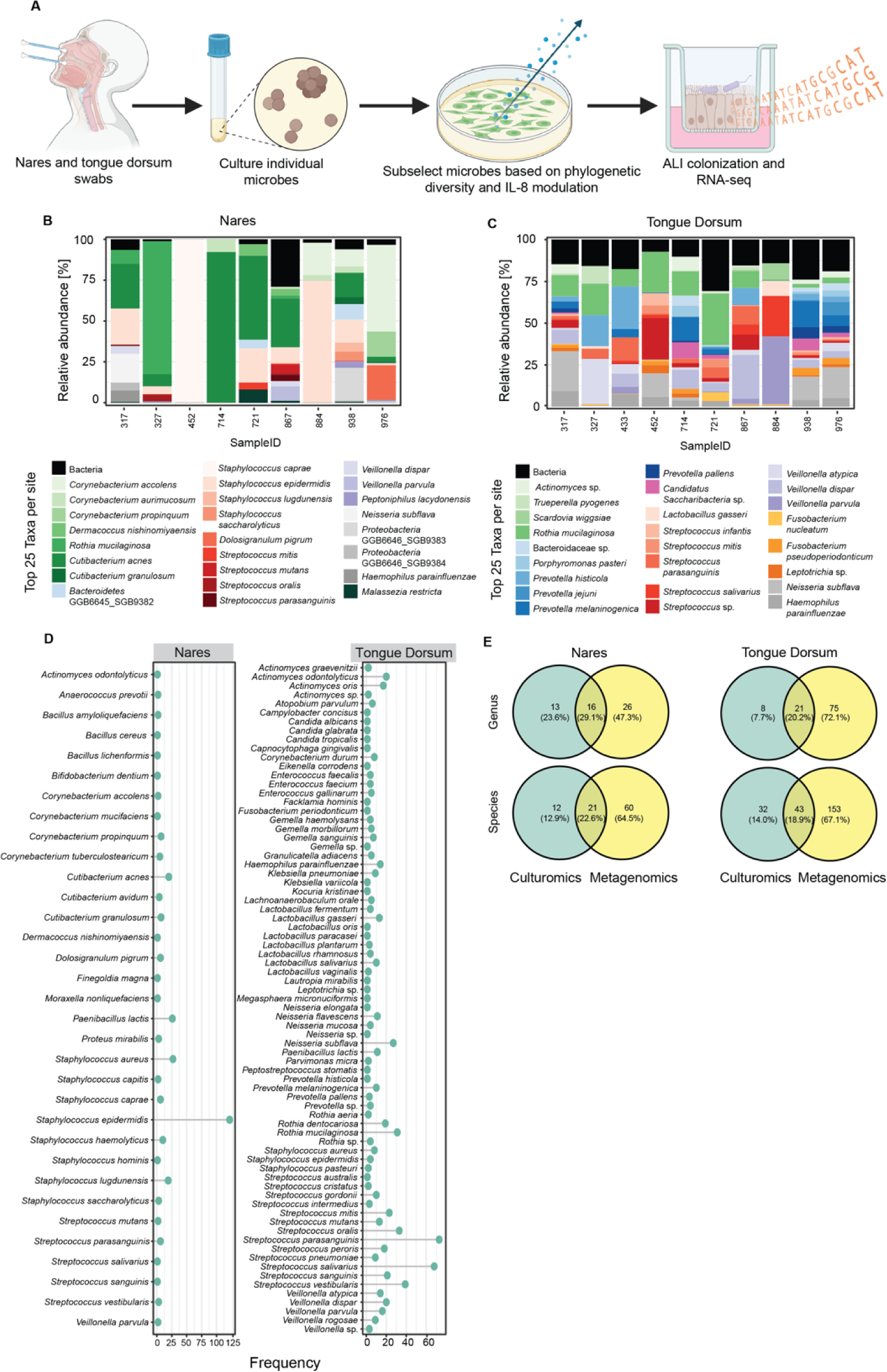
Study design and nares and tongue dorsum microbiome characterization. **A**) Schematic of microbial isolation and study design. The nares and tongue dorsum of lung cancer patients were swabbed, and individual microbes were isolated through culturomics. Filtered microbial supernatants were applied to A549 lung epithelial cells, and changes in IL-8 secretion was quantified. Microbes were subselected based on phylogenetic diversity and modulation of IL-8 and used to colonize ALIs for RNA-seq analysis. **B**) Relative abundance plots of the most abundant bacteria and fungi are shown for select samples from the nares and tongue dorsum that were used for cultivation. **C**) Total frequency for which each species was detected using different culturomics conditions per site. **D**) Overlap of genera and species recovered by culturomics compared to metagenomics for the two sites sampled. In the Venn diagram, the number of unique isolates and the percent of total unique isolates isolated using culturomics only, metagenomics only, and both is shown.

Following metagenomic analyses, we cultured 1,027 microbes as previously described^28^ from the samples from 11 randomly selected patients (Fig. 1A). Our goal was to isolate a diversity of microbial species for our ALI host-interaction screen. First, we measured the frequency that each microbial species was isolated (Fig. 1D). As anticipated, common microbiota such as *S. epidermidis* and *Streptococcus* spp. were isolated with greater frequency, allowing potential investigation of strain-specific effects. Most species were cultured fewer than 20 times in both the oral and nasal samples, likely demonstrating patient-specific effects. To examine how representative cultivars were of the predicted metagenome, we compared the number of uniquely identified microbes using metagenomics versus those identified using culturomics at both a genus and species level (Fig. 1E). Unsurprisingly, we saw more unique genera/species identified using metagenomics than using culturomics at both sites. While metagenomics identified 76-91% of the total genera/species detected by either method, culturomics only identified 28-53%, with oral samples tending to recover fewer genera/species than the nasal samples.

### Microbial selection and colonization on ALIs

We then sought to rationally subset the 1,027 isolated respiratory microbes for follow-up investigations in ALIs, and at the same time gain insights into the diversity of potential phenotypes of interest generated by different phylogenies of microbes. Because secreted microbial metabolites are immunogenic and microbial immunogenicity can be species- and strain-specific, we first examined the inflammatory potential of microbial metabolites on 2D lung cell cultures as a high throughput means of screening the isolated microbes. We collected conditioned bacterial supernatant from each isolated microbe, applied the supernatants (20% v/v) to A549 lung epithelial cells overnight, and measured the concentration of six human inflammatory cytokines (IL-8, IL-6, IL-1, IL-12, IL-10, and TNF) using cytokine bead array (CBA). Interestingly, bacterial conditioned supernatant only caused significant changes to the secretion of IL-8, a neutrophil chemoattractant^29^. Notably, modulation of IL-8 was not coupled to phylogeny (Fig. S1A). At the genus level, the median log2 fold change for nearly every genus was decreased IL-8 secretion but within-genus and even to the strain level, there was notable isolate-specificity with a large log_2_ fold change (FC) range in IL-8 secretion (Fig. S1B).

We ultimately selected 56 microbes for colonization of ALIs, prioritizing based on genetic diversity (different species) and maximizing the effect on IL-8 secretion (increased or decreased secretion) (Fig. 2A). We also included a GFP-tagged *S. epidermidis* strain Tü3298^30^ for a visualization and colonization control, a skin-derived *S*. *epidermidis* strain 3F3^31^ as an additional colonization control, and water as a vehicle negative control. 10^7^ CFUs of each microbe (colony forming units, a measure of live bacteria) or vehicle were applied to the apical surface of mature tracheobronchial ALIs for 18 or 48 hours (Fig. 2B) to approximate spatial aspects of physiological conditions. At each time point, total eukaryotic RNA was extracted from the tissue and processed for RNA-seq, basal media was collected for IL-8 quantification, and the apical surface was washed to quantify CFUs to approximate bacterial growth/load (Fig. S2). We observed several growth patterns: 1) a decline in CFUs at 18 hours followed by increasing CFUs by 48 hours, 2) continuous bacterial growth at both timepoints, 3) a continuous decrease in CFUs at both timepoints, or 4) CFUs at a steady state, and these patterns were genus, species, and strain-specific. For example, most *Staphylococcus* spp. experienced an initial decline followed by growth equilibrating at ∼10^7^ CFUs by 48h, though the magnitude of the decline and ultimate equilibration differed by species (e.g., *S*. *haemolyticus’* and *S*. *lugdunensis’* vs. *S. aureus).* Similarly, *Streptococccus* spp. equilibrated around ∼10^5^ CFUs with varying trajectories, with some isolates (*Str*. *parasanguinis*, *salivarius*, and *vestibularis*) decreasing through 48 hours. We note that isolates may differ in their adhesive properties, thus this analysis is at best an approximation of microbial growth and load.

**Figure 2:**
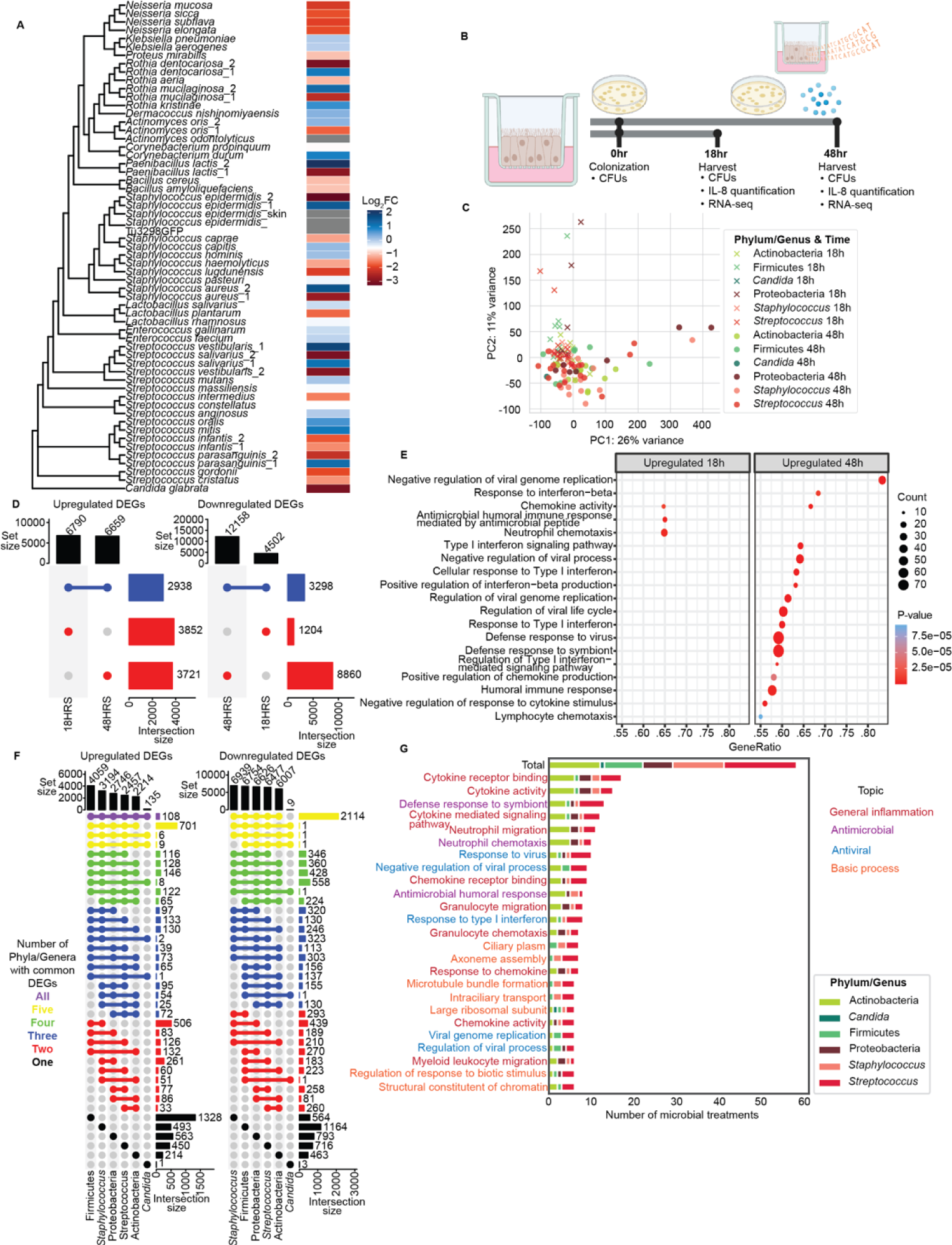
Microbial colonization of epithelium results in gene expression changes, particularly in antiviral immunity, that are not phylogenetically conserved. **A**) Phylogenetic tree of microbes used to colonize ALIs. Heatmap shows the log_2_ fold change (FC) of A549 lung epithelial cell IL-8 secretion following exposure to different microbial supernatants. **B**) Schematic of the RNA-seq experiment. ALIs were colonized with 10^7^ microbes or vehicle (H_2_O), at which point 0 hour (dosing) CFUs were quantified by dilution plating. ALIs were colonized for 18 and 48 hours (n=3-5). At each harvest, CFUs were quantified from an apical wash, IL-8 in the basal media was quantified, and total RNA of the epithelial cells was sequenced. **C**) PCA plot of genome-wide transcriptional responses of ALIs colonized for 18 (“x”) or 48 hours (“o”) relative to vehicle control. Dots are colored based on phylum or select genera. **D**) Upset plot of cumulative significantly (adjusted P-value < 0.1) differentially expressed genes (DEGs) of ALIs separated by timepoint, 18 or 48 hours. Set size indicates total number of up- or downregulated DEGs per timepoint. X-axis (timepoint) is sorted by decreasing set size. Colored dots indicate the timepoints included in the intersection size (number of DEGs). Lines connect the colored dots, when the DEGs are shared between both timepoints. DEGs that are shared between both timepoints are blue and DEGs unique to one timepoint are red. **E**) Gene set enrichment analysis of cumulative upregulated DEGs at 18 and/or 48 hours. Top 20 significantly upregulated gene sets (FDR adjusted P-value < 0.05) with the highest gene ratio (percent of genes within the gene set that are significantly upregulated). The top 20 gene sets are plotted by time if the gene sets are significantly upregulated at 18 or 48 hours. The size of the dot represents the number of enriched core genes in that particular gene set. The color of the dot represents the FDR adjusted P-value. **F**) Upset plot of significantly (adjusted P-value < 0.1) cumulative up- and downregulated DEGs after colonization for 48 hours separated by phylum/genus. Set size represents total number of up- or downregulated DEGs. X-axis (phylum/genus) is sorted by decreasing set size. Colored dots indicate the phyla/genera used to calculate the intersection size (number of DEGs). Lines connect phyla/genera when the DEGs are shared between multiple categories. Plot is colored based on the number of phyla/genera the DEGs are shared between. **G**) Gene set enrichment analysis of statistically significant (adjusted P-value < 0.05) upregulated DEGs at 48 hours, separated by phylum/genus. Top 20 gene sets that were significantly upregulated by the largest number of microbial treatments. The x-axis represents the number of microbial treatments that were significantly upregulated in each gene set. Gene sets are colored based on general function. “Total” gene set shows the total number of microbial treatments per phylum/genus category. For each gene set, the bar is then colored based on the number of microbial treatments, for each category, that significantly upregulated for that gene set.

### Microbial modulation of host gene expression is not phylogenetically conserved

First, we sought to determine if bacterial growth/load was the primary driver of the transcriptional changes observed, as it is conceivable that excessive growth on the ALIs would trigger a broad epithelial response. We used a linear mixed effects model (MaAsLin2^32^) to determine if CFUs were a confounder for differential gene expression. We observed that only 68 genes (of 42,609 genes expressed in our model) were driven by CFUs, none of which were in our pathways of interest for subsequent analyses.

Next, we examined global differences in the effects of microbial colonization on host gene expression as a function of time. A qualitative comparison of 18 and 48 hour timepoints with principal components analysis (PCA) showed that most of the microbial treatments at both 18 and 48 hours clustered together and that the outliers were not phylogenetically similar (Fig. 2C). This suggested that species-specific epithelial responses to colonization could be primarily compartmentalized to certain pathways such as innate immune response or barrier integrity. Differentially expressed genes (DEGs) at each time point were similar, with substantial overlap between timepoints, though there were markedly more downregulated genes at 48 hours (Fig. 2D). Examining the top 20 gene sets with the highest gene ratio (percent of genes enriched in the gene set) from either time point, we observed that the overwhelming number of enriched gene sets were from the 48 hour time point, with low gene ratios at 18 hours (Fig. 2E). Surprisingly, we saw enrichment in several antiviral gene sets, including Type 1 interferon (IFN-I), which is a canonical antiviral response, as few studies have identified or investigated bacterial stimulation of interferon. As these results suggested that transcriptional differences accumulate and are most discernable at 48 hours, we focused on data from the 48 hour timepoint for the remainder of analyses.

To investigate if the effects on epithelial gene expression at 48 hours were phylogenetically driven, we compared the number of upregulated and downregulated DEGs between different microbial treatments (Fig. 2F), categorizing largely by phyla, with genus-level designations *Staphylococcus (S)*. and *Streptococcus (Str.)* spp. given their predominance and *Candida* given there was only one isolate. Interestingly, the number of DEGs did not correlate with the group size as one might anticipate; for example, though *Streptococcus* had the largest number of treatments, its effect on the epithelium was moderate, as measured by the total number of upregulated DEGs. Strikingly, *Streptococcus,* despite having a comparable number of unique upregulated DEGs, shared few upregulated DEGs with *Staphylococcus* and other Firmicutes. *Staphylococcus* shared more upregulated DEGs with Proteobacteria than *Streptococcus* did with *Staphylococcus* or other Firmicutes. Regardless of how many groups compared, Actinobacteria shared few upregulated DEGs and *Candida* even fewer. The strong similarity between *Staphylococcus* upregulated DEGs and those of other Firmicutes is unsurprising, given the closer phylogeny. However, *Streptococcus’* and Actinobacteria’s few upregulated DEGs shared with other groups suggests that microbial induction of host epithelial genes is not highly phylogenetically conserved and can vary in both effect and quantity. Contrastingly, with the exception of the fungus *Candida,* which only included one treatment and is phylogenetically most removed from the bacterial isolates, there were more shared downregulated DEGs, indicating that aspects of microbial inhibition of host epithelial genes are likely conserved.

We then examined if phylogenetically more similar microbes induced similar gene expression changes in the 25 gene sets that were enriched in the most microbial treatments. Most of the enriched gene sets, irrespective of phylogeny, were related to general inflammation, such as neutrophil migration/chemotaxis and cytokine binding/activity. Other gene sets included antiviral response (response to virus and IFN-I). Notably, these enrichments crossed phylogenetic lines – i.e., almost all gene sets were upregulated by at least one member of each phylogenetic group – but surprisingly, no gene sets were upregulated by all members of a phylogenetic group. This further supports the notion that microbial induction of host gene expression is not phylogenetically conserved.

### Microbial induction of interferon is not phylogenetically determined

We then further investigated the surprising enrichment in IFN-I gene sets, because IFN-I (as well as Type 3 interferon (IFN-III), which is also expressed in the respiratory epithelium) is canonically a strictly antiviral immune response that results in the activation of a number of interferon-stimulated genes (ISGs). To quantitate bacterial stimulation of interferon, we manually curated a list of 61 ISGs from the literature^33^ representing IFN-I and IFN-III signaling, which have almost identical signaling cascades and ISGs, by selecting responsive genes using hierarchical clustering. Interferon stimulation at 48 hours was strikingly dichotomous (Fig. 3A) with approximately half of the microbes strongly stimulating interferon (i.e., strongly upregulating ISGs), while the other half had a minimal effect. For each microbial treatment, we calculated the median log_2_FC for the ISGs to create an ISG score (Fig. 3A), which we confirmed using previously published 6-^34^ and 30-gene^35^ ISG scores (Fig. S3). Based on the hierarchical clustering and ISG score, we then classified the microbes as ISG non-stimulator versus simulators, characterized by ISG score as well as a significant difference in average log_2_FC of ISGs (Fig. 3B, P=9.9e-11, two-sided Mann-Whitney U test with Bonferroni correction). We compared the effect of microbial colonization on interferon genes to determine whether the microbes were inducing IFN-I or IFN-II; however, we saw no significant differences.

**Figure 3:**
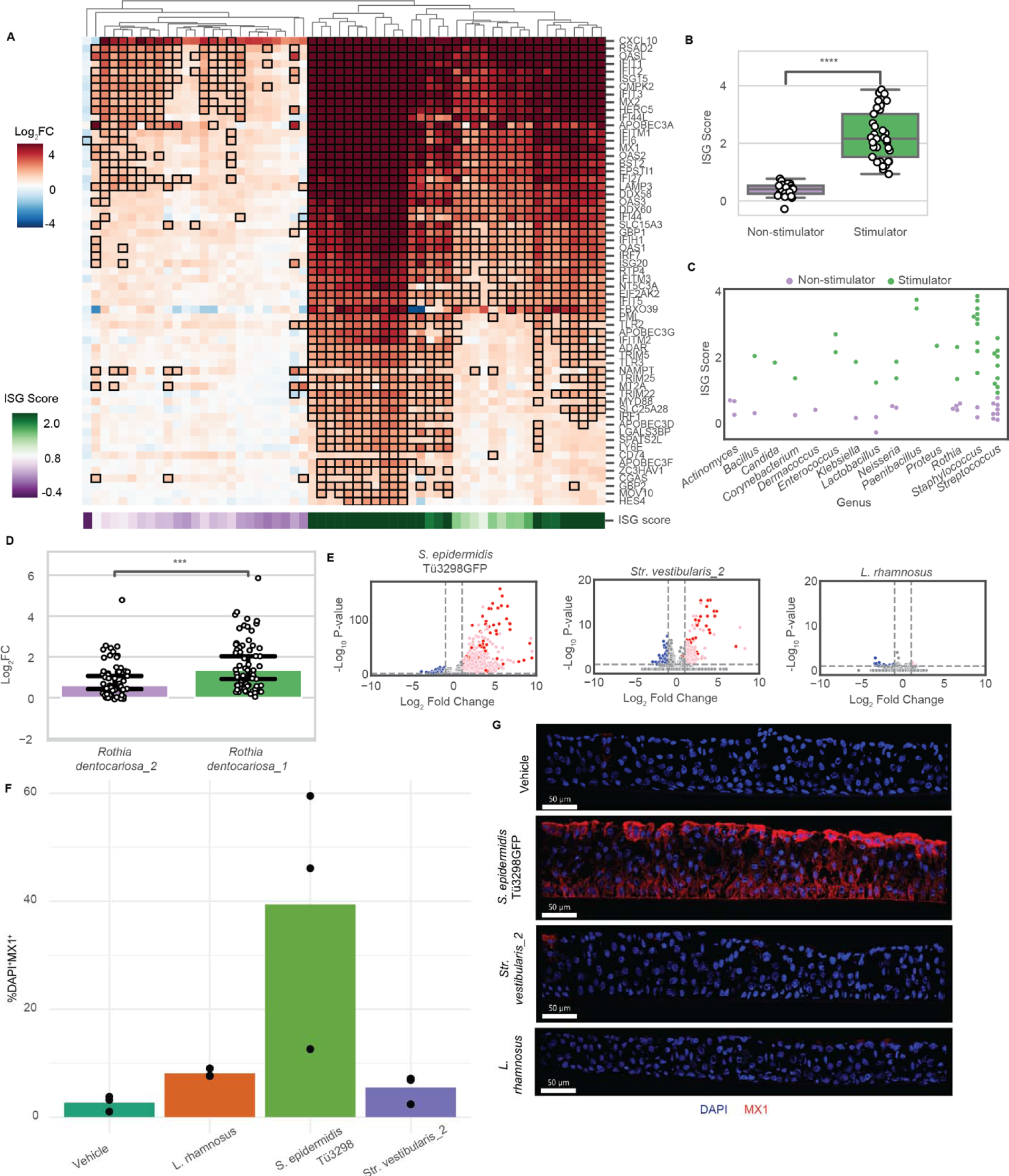
Species and strain level differences in microbial induction of antiviral interferon pathways. **A**) Heatmap (log_2_ fold change (FC)) of the transcriptional response of manually curated interferon-stimulated genes (ISGs) in ALIs colonized with microbes for 48 hours relative to vehicle control. Each column represents a different microbial treatment and each row a different ISG. Cells were outlined to indicate statistical significance (FDR adjusted P-value < 0.1). Microbial treatments were clustered into interferon non-stimulator and stimulator categories based on hierarchical clustering. ISG score was determined based on the median log_2_FC of the ISGs. Below the heatmap is each microbial treatment’s ISG score; green represents higher log_2_FC (stronger interferon induction) and purple represents lower log_2_FC (less interferon induction). **B**) Boxplot of ISG score of ISG non-stimulator (purple) vs. stimulator (green). Each point represents a microbial treatment; box edges indicate 25^th^ and 75^th^ percentiles. **C**) Dot plot of each microbe’s ISG score where microbial treatments are sorted by genus. Dots are colored purple to represent interferon non-stimulators and green to represent stimulators. **D**) Bar graph of the log_2_FC for each ISG with each of the two *Rothia dentocariosa* strains, one of which is an interferon non-stimulator (purple) and the other an interferon stimulator (green). **E**) Volcano plots of DEGs for the three microbial treatments used for validation of transcriptional results by immunofluorescence. Genes (points) colored in blue are significantly downregulated (adjusted P-value < 0.1), pink are significantly upregulated, and red are significantly upregulated ISGs. **F-G**) Immunofluorescence of ALI colonized for 48 hours with a microbe or vehicle (n=3). **F**) Quantification of MX1 by histocytometry represented as bar graphs representing the percentages of DAPI^+^MX1^+^ cells per condition. Data shown are representative of 3 technical replicates per condition with at least 2 ALI sections per replicate. Quantification of the percent of cells that express *MX1*. Each data point represents the average of three sections. **G**) Representative immunofluorescent sections (8 µm) of ALI cultures after 48 hr of bacterial/vehicle exposure stained for nuclei (DAPI), and MX1 (red) to reveal downstream of IFN response. Scale bar 50 μm, in white on the left corner. For relevant plots, statistical analysis is two-sided Mann-Whitney U test with Bonferroni correction. * indicates P-value <0.1, **<0.05, ***<0.01, ****<0.001.

Strikingly, the ability of a microbe to stimulate ISG was not predicted by phylogeny, even at the strain level. Nearly every genus included both non-stimulators and stimulators with a wide range of stimulatory potential (Fig. 3C). Interestingly, we even observed strain-level differences, e.g., *Rothia dentocariosa* (Fig. 3D). Strains could moreover differ temporally in their ability to activate interferon; a *Paenibacillus lactis* strain and a *Str*. *parasanguinis* strain induced interferon at 18 hours while the other strains of those species (and the vast majority of ISG stimulators) induced interferon only by 48 hours (Fig. S4). Finally, to confirm that bacterial growth/load did not solely explain the difference in interferon stimulation, we compared CFUs at 48 hours between the ISG non-stimulator and stimulator microbial treatments and found no significance (Fig. S5, P=0.12, two-sided Mann-Whitney U test).

To experimentally validate interferon activation, we selected three microbial treatments (*Str. vestibularis*, *L*. *rhamnosus*, and GFP *S*. *epidermidis* Tü3298) and validated MX1 expression, a direct antiviral mediator protein known to be induced by interferon, by immunofluorescence. These three isolates represented the extremes of transcriptional effects on colonization; *S*. *epidermidis* elicited the most upregulation, *Str*. *vestibularis* had the most downregulated genes, and *L*. *rhamnosus* had a surprisingly minimal effect on any genes despite robust growth on ALIs (Fig. 3E). Immunofluorescence quantifying the percent of cells expressing MX1 confirmed increased expression of MX1 in ALIs colonized with *S*. *epidermidis*, while minimal changes in MX1 expression was observed for the other two treatments (Fig. 3F-G). While *S*. *epidermidis*’ and *L*. *rhamnosus*’ effect on MX1 expression matched their respective stimulator and non-stimulator category, *Str*. *vestibularis’* lack of MX1 expression was discordant with the RNA-seq results. One explanation is that while *Str*. *vestibularis* can induce gene expression of *MX1*, expression does not pass the threshold for protein expression. Alternatively, *Str*. *vestibularis*’ interactions with the ALIs may lack secondary interactions necessary for MX1 protein expression (or the inverse). Taken together, the degree of species and strain specificity in microbial induction of interferon suggests that small variations or changes to the respiratory microbiome may have significant effects on antiviral immunity.

### Microbial induction of interferon is independent of antibacterial immunity and epithelial barrier

Host innate immunity includes a wide array of antibacterial immune responses to resist genetically diverse bacteria. Our 58 microbes included a mix of canonical commensals and opportunistic pathogens, and thus we anticipated that some microbes would be more immunogenic while others may be immunosuppressive. To investigate if our microbial treatments differentially activated those antibacterial immune responses, we manually curated a list of 210 genes including non-interferon cytokines and antimicrobial peptides from MSigDB^36^ with gene ontologies^37,38^ relating to cytokines and antimicrobial immune response, which we termed antibacterial innate immune response. Of those antibacterial innate immune genes, 36 of the genes differentially responded to microbial treatment (Fig. 4A). An antibacterial score was generated based on median log_2_FC, identifying two classes of microbial induction: weak versus strong stimulator, again identified by hierarchical clustering and having significantly different antibacterial scores (Fig. 4B). Like interferon response, microbial phylogeny did not predict whether the microbe would be a strong or weak effector of antibacterial immunity (Fig. 4C). However, unlike interferon, induction was not dichotomous and was instead more continuous within and between microbial clades. To experimentally validate these broad categorizations, we measured the concentration of IL-8 (*CXCL8*) secreted into the basal media following microbial colonization for 18 and 48 hours. Nearly all microbial treatments slightly increased secretion of IL-8 at 18 hours and significantly increased secretion after 48 hours (Fig. 4D), and secretion was correlated with transcriptional levels (Fig. S6, ρ=0.72, P<2.2e-16, Spearman’s correlation coefficient). Unlike interferon, when we compared CFUs at 48 hours between strong and weak antibacterial inducers, we found a significant difference (Fig. S7, P=4.2e-06, two-sided Mann-Whitney U test). While none of the individual antibacterial genes were predominantly driven by CFUs per our multivariate model, grouping these genes together increased the effect to be statistically significant. This was in striking contrast to interferon’s induction, which was independent of microbial load.

**Figure 4:**
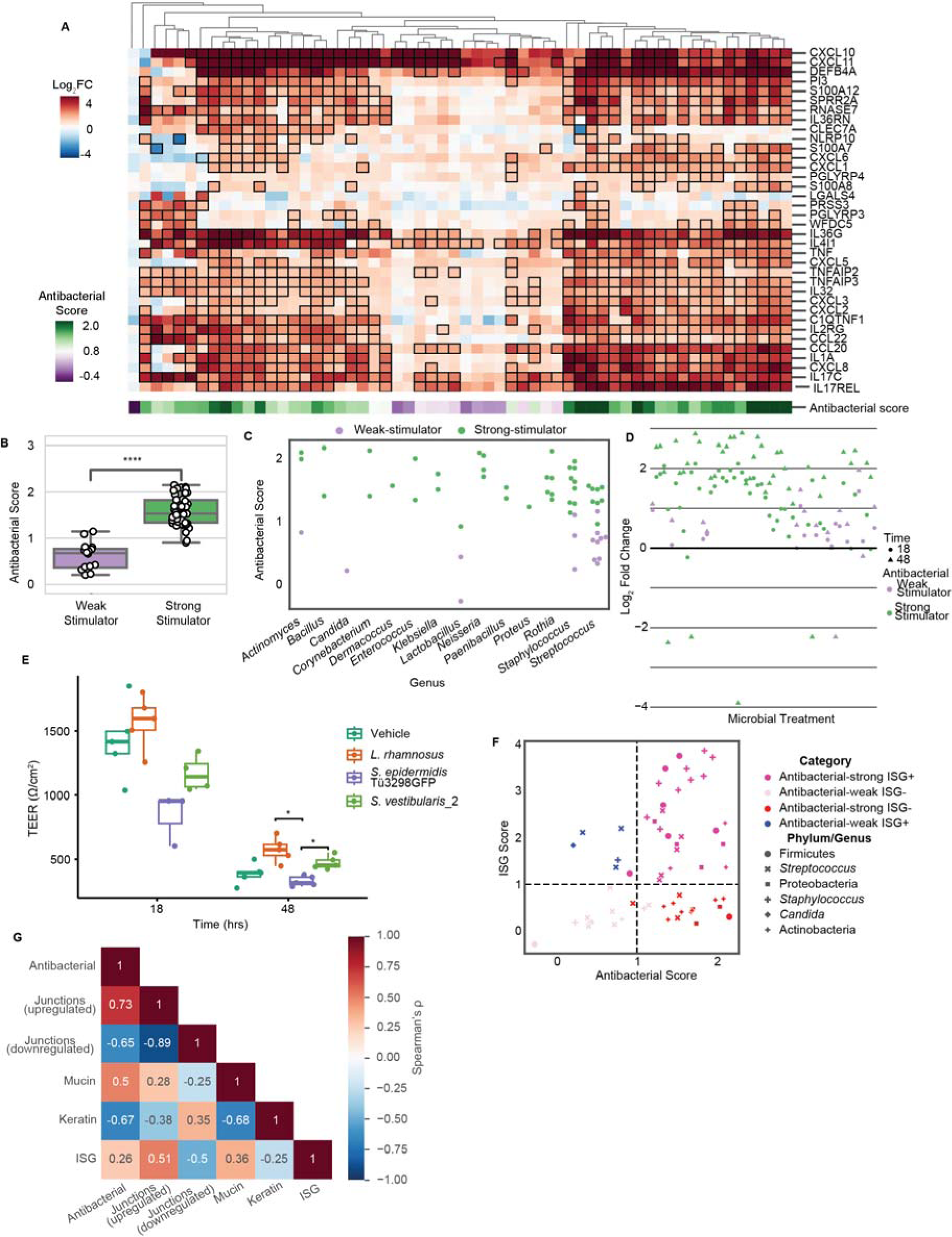
Microbial modulation of antibacterial genes are correlated with barrier genes while uncoupled with ISG expression. **A**) Heatmap, as in Fig. 3A visualizing the transcription response of manually curated antibacterial genes (non-interferon cytokines and antimicrobial peptides) in ALIs colonized with microbes for 48 hours relative to vehicle. Black boxes indicate statistical significance (FDR adjusted P-value < 0.1). Microbes were grouped into antibacterial strong and weak stimulators based on hierarchical clustering. Below the heatmap is the microbe’s antibacterial score, which was calculated from the median log_2_FC of the curated genes. **B**) Boxplot of each microbe’s antibacterial score, grouped based on the microbe’s antibacterial weak versus strong stimulator category. Box edges indicate 25^th^ and 75^th^ percentiles and statistical analysis is two-sided Mann-Whitney U test with Bonferroni correction. **C**) Dot plot of each microbe’s antibacterial score separated by genus and colored by antibacterial weak stimulator (purple) or strong stimulator (green) category. **D**) Log_2_FC in basal IL-8 secretion relative to vehicle (n=3). Each point represents a different microbial treatment, shaped as a circle for 18 hour timepoint or triangle for the 48 hour timepoint. Points are colored based on their antimicrobial weak versus strong stimulator category. **E**) Boxplot of transepithelial electrical resistance (TEER) following microbial colonization or vehicle for 18 or 48 hours (n=3-5). Statistical analysis is ANOVA with post-hoc Tukey test. **F**) Each microbial treatment is plotted based on its ISG (y-axis) and antibacterial (x-axis) scores, colored based on its ISG and antibacterial stimulation category and shaped based on its phylum/genus. **G**) Correlation matrix between the median log_2_fold change of antibacterial, upregulated junctions, downregulated junctions, mucin, keratin, and ISG gene lists. Each box contains the Spearman’s correlation coefficient (ρ) and is colored based on the P-value. For all relevant plots, * indicates P-value <0.1, **<0.05, ***<0.01, ****<0.001.

Barrier integrity affects the penetrance of microbes and microbial products and thus immunogenicity. Some components of the epithelial barrier, such as mucus secretion, are often modulated by microbial colonization^1^ while other components, such as tight junctions, are common targets of microbial pathogens^39^. Based on this, we also examined epithelial barrier genes. We curated a gene set of keratin, mucin, and junction (inclusive of tight junctions, gap junctions, and adherens junctions) genes (Fig. S8), similar to ISGs and antibacterial gene lists, from the literature^40,41^ and HGNG (HUGO Gene Nomenclature Committee)^42^. We experimentally validated a subset of the transcriptional changes observed in the epithelial barrier genes. We measured the transepithelial electrical resistance (TEER), a measurement of barrier permeability, of ALIs treated with GFP *S*. *epidermidis* Tü3298, *Str*. *vestibularis*, and *L*. *rhamnosus* at 18 hours and 48 hours (Fig. 4E). *L*. *rhamnosus*, which was an ISG and antibacterial non-stimulator, showed increased TEER and therefore decreased permeability, while *S*. *epidermidis*, an ISG stimulator and antibacterial strong stimulator, decreased TEER/increased permeability. *Str. vestibularis*, an ISG stimulator and weak antibacterial stimulator, did not have a significant effect on TEER - altogether, these results aligned with each microbe’s degree of antimicrobial stimulation and not ISG stimulation. This suggests there may be a connection between microbial induction of antibacterial pathways and epithelial barrier genes.

To further generalize these potential links between epithelial barrier, antibacterial immunity, and ISG induction, we determined if the same microbial species were inducing interferon and antibacterial immunity, and therefore, were more generally immunogenic. Notably, there was little overlap in a microbe’s ability to induce interferon versus antibacterial immunity; some microbes induced both, one of the two, or neither, again with no discernable phylogenetic explanation (Fig. 4F). In addition, ISG expression did not have a strong correlation with mucin or keratin gene expression. However, ISG expression did strongly correlate with junction genes. Furthermore, antimicrobial immunity gene expression was strongly negatively correlated with keratin gene expression and strongly positively correlated with mucin gene expression (Fig. 4G, Spearman’s correlation coefficient). The weak correlation between mucin/keratin genes and junction genes suggests that they are regulated separately. Taken together, we concluded that microbial induction of host interferon is independent from modulation of host antimicrobial immune response and most components of the epithelial barrier, which are more closely coupled in the respiratory epithelium.

### In silico identification of potential immunomodulatory microbial pathways

A recurring theme in our examination of epithelial response to microbial colonization was that stimulation of different host pathways was largely independent of phylogeny. It would then follow that there could be numerous mechanisms by which microbes can stimulate pathways such as interferon. We conducted an *in silico* analysis of the microbial genomes used in this study to identify potential microbial genes/pathways that may stimulate host interferon. While we expected microbial induction of interferon to be poorly conserved, given the lack of phylogenetic similarity between ISG stimulators, we hypothesized that some broad pathways may be conserved.

We quantified the presence or absence of broad functional categories of virulence factors (via VFDB^43^) in each genome: adherence, invasion, effector delivery system, motility, exotoxin, exoenzyme, immune modulation, biofilm, nutritional/metabolic factor, stress survival, post-translational modification, antimicrobial activity/competitive advantage, regulation, and others (Fig. 5A). Overall, microbes were most likely to have virulence factors related to nutritional/metabolic function or immune modulation. When hierarchically clustered, microbes did not cluster strictly phylogenetically. More importantly, interferon non-stimulators and stimulators did not cluster separately. This suggests that overall patterns of virulence factors do not contribute to a microbe’s ability to induce interferon.

**Figure 5.**
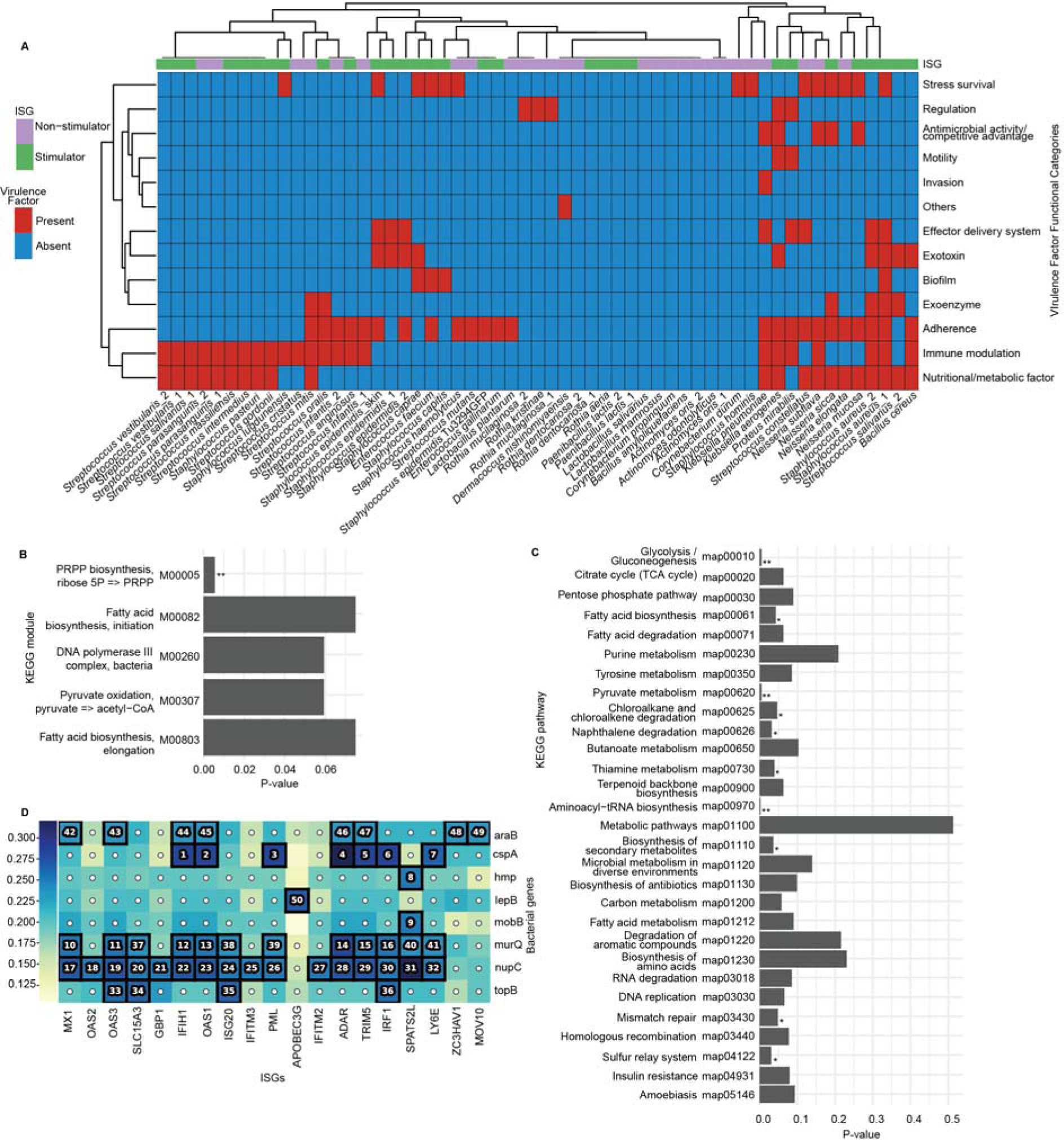
In silico prediction of potential microbial pathways associated with interferon induction. **A)** Heatmap of broad virulence factor functional category presence/absence for each microbial genome annotated with VFDB. Red indicates that the microbial genome contained at least one virulence factor in that category and blue indicates there were no virulence factors identified in that category. Above the heatmap, each microbial treatment is annotated by its ISG non-stimulator vs stimulator ability (purple represents non-stimulator and green represents stimulator). The heatmap’s x and y axes are hierarchically clustered. **B-C)** All KEGG modules/pathways enriched, with KEGG orthologs that were differentially abundant between ISG non-stimulators and stimulators. For each enriched module/pathway, the P-value for the differential abundance between ISG non-stimulators and stimulators is displayed on the x-axis (Mann-Whitney U test). **B)** KEGG modules. **C)** KEGG pathways. **D)** Correlation matrix (NMI, normalized mutual information) between host ISG log_2_FC and microbial gene presence. The heatmap is colored by correlation (blue is high correlation and white is low correlation). Correlations are outlined if statistically significant (FDR adjusted P-value < 0.05). The number in the outlined correlations indicates the rank of the correlation ordered by decreasing association strength.

Next we used FMAP^44^ to determine differentially abundant KEGG orthologs between ISG non-stimulator and stimulator microbes and identified KEGG modules (Fig. 5F) and pathways (Fig. 5G) that were enriched for the differentially abundant orthologs. Among the KEGG modules enriched in differential orthologs, the only module significantly differential between ISG non-stimulators and stimulators was the conversion of ribose 5-phosphate to phosphoribosyl pyrophosphate (PRPP). PRPP is used in the biosynthesis of nucleotides, aromatic amino acids, and cofactors like NAD, among other biomolecules^45^. Significantly differential KEGG pathways included biosynthesis of secondary metabolites. Secondary metabolites are bioactive molecules that are not required for microbial survival but confer a competitive advantage and can often modulate human immunity^46^.

Finally, we used HAllA^47^, a dimensionality reduction algorithm that could accommodate the large number of microbial genes, to find associations between microbial genes and ISG log_2_FC (Fig. 5H). Out of the genes that were significantly associated with several ISGs, *araB*, which converts ribulose into ribulose 5-phosphate and vice versa^48^, was particularly interesting because ribulose 5-phosphate can be converted into ribose 5-phosphate^49^ (see discussion). Taken together, the variability in bacterial pathways associated with ISG stimulation emphasizes the diversity of potential interaction mechanisms with the epithelium, with the exception of the potentially more conserved pentose phosphate pathway.

## Discussion

Here, we presented a broad analysis of how microbial phylogenetic diversity affects host-microbe interactions in the respiratory tract. Using a complex 3D epithelial cell culture model, we characterized the effect of colonization with one of 57 phylogenetically diverse bacteria and one fungal isolate on epithelial gene expression.

A notable finding was a broad, dichotomous ability of phylogenetically diverse microbes to stimulate host interferon. Differential microbial induction of interferon has implications for antiviral immunity/susceptibility and microbiome-based therapies for respiratory viral infections. IFN-I and IFN-III, the two types of interferon present in epithelium, are critical for antiviral immunity^7^ and thus, have predominantly been studied in that context. The few studies that have investigated bacterial induction of interferon focused on peripheral blood mononuclear cells, especially dendritic cells and monocytes^51–57^. However, we found that half of our microbial treatments (bacterial and fungal) induced epithelial interferon following 48 hours of colonization. Interestingly, the *Candida glabrata* isolate was an interferon stimulator, which to the best of our knowledge is the first demonstration of fungal induction of interferon. We note that while we categorized our microbes as interferon non-stimulators versus stimulators, it is likely that this activation is not entirely binary. A few microbes were able to induce interferon by 18 hours – in a mechanism unrelated to microbial growth rate – and all stimulator microbes by 48h, it is possible that there are also differences in how quickly a microbe can induce interferon, which has clinical relevance in the ability and rapidity of the microbiome to modulate antiviral immunity. Another striking finding was that a microbe’s potential to stimulate interferon was not phylogenetically driven; we observed species- and strain-level differences, which aligns with other studies^51–54^. Furthermore, a microbe’s broad categorization as a typical commensal versus pathogen also did not correspond with their ability to induce interferon. For example, both *S*. *aureus* isolates, common respiratory pathogens, were interferon stimulators while *Klebsiella pneumoniae*, another major respiratory pathogen, was a non-stimulator.

In addition, we also observed that a microbe’s potential to stimulate interferon was not dependent upon its potential to stimulate antibacterial immunity, i.e., a microbe that was an interferon stimulator was not necessarily an antibacterial stimulator, and vice versa. This was consistent with another study which found that bacteria that induced IFN-I interferon were poor NF-κB inducers, and vice versa^55^. Induction of antibacterial immunity was moreover driven in part by microbial load, which was unsurprising in that more microbes led to more stimulation of pattern recognition receptors and consequently, an elevated immune response. In contrast, interferon induction did not correspond to microbial load – microbes that grew exponentially and microbes that consistently appeared to die or not grow on the ALIs both induced interferon. Together, this leads us to believe that microbes can induce host interferon through different mechanisms than used to induce antibacterial immunity.

Our results are notable because there has traditionally been a divide between bacterial pattern recognition receptors, which recognize extracellular microbes and initiate antibacterial immune responses, and viral pattern recognition receptors, which targets intracellular microbes and induced antiviral immune responses^58^. Interferon, as an antiviral immune response, has canonically been associated with intracellular viral pattern recognition receptors. Therefore, microbial induction of epithelial interferon may be a previously uncharacterized function of traditional antibacterial immune response, a previously uncharacterized means of stimulating viral pattern recognition receptors, or an uncharacterized mechanism. Several different mechanisms of bacterial induction of interferon have been demonstrated. For example, outer membrane polysaccharide A has been shown to induce IFN-I through TLR4^54^. Bacterial DNA can induce IFN-I through TLR9^56^, and bacterial RNA can enhance IFN-I through TLR3, TLR9^52^, cGAS/STING, and RIGI-like receptors^55^. Secreted metabolites such as desaminotyrosine (DAT)^59^ and secondary bile acids^57^ also have been shown to promote IFN-I signaling. However, all of these mechanisms were demonstrated through microbial interactions with phagocytotic immune cells and may not translate to epithelial cells. For example, microbial nucleotide stimulation of TLR3, TLR9, cGAS/STING, and RIGI-like receptors is proposed to occur following phagocytosis of microbes. However, this could not occur in epithelium. Furthermore, given that interferon stimulators were not necessarily phylogenetically similar, it is plausible that there are many mechanisms through which microbes induce interferon and that those mechanisms are not necessarily conserved.

While the microbes we used are phylogenetically dissimilar, and we expected multiple potential mechanisms of microbial induction of interferon, one trend we found was an association between the pentose phosphate pathway and interferon induction. The pentose phosphate pathway, through ribose 5-phosphate and PRPP, can synthesize purine and pyrimidine nucleotides and aromatic amino acids including tryptophan^45^. PRPP’s potential to synthesize nucleotides is interesting in the context of previous studies that demonstrated that phagocytosed bacterial nucleotides induced interferon^56,52,55^. However, PRPP can also be converted into a variety of other bioactive molecules, which could modulate bacterial or host gene expression^45^.

In conclusion, we demonstrated that microbial induction of interferon is highly heterogenous, not phylogenetically driven, and occurs independently of microbial induction of more general inflammatory processes. This dataset builds a foundation for understanding how respiratory microbiome biodiversity affects microbe-epithelial interactions, particularly in the context of antiviral innate immunity.

## Materials and Methods

### Lung cancer patient recruitment

We recruited 47 patients (46-83 years old) with established evidence of adenocarcinoma lung cancer. Written informed consent was obtained from all participants and the study protocol was approved by the Human Subjects Institutional Review Board at Hartford Hospital, Hartford, USA and performed in the context of U19AI142733 grant at the Jackson Laboratory.

### Sample collection

Nares and oral samples were obtained using sterile swabs (Puritan™ PurFlock Ultra, #22-029-506, Guilford, ME, USA). For nares, two swabs pre-moistened in nuclease-free water (Qiagen, Hilden, Germany) were inserted 2 cm into one nostril and rotated against the anterior nasal mucosa for up to 10 seconds. The tongue dorsum was gently rubbed with two dry swabs for up to 30 seconds.

One swab each was stored in 350 μl Tissue and Cell lysis solution (Lucigen, #MTC096H, Middleton, WI) lysis buffer with 100 μl glass beads (0.1 diameter, BioSpec Products, Cat No. 11079101, Bartlesville, OK, USA). The other swab was placed and stored in R2A (Innovation Diagnostics, LAB203-A) for future microorganism recovery. All samples were stored at −80°C until shipping. Samples were shipped on dry ice to The Jackson Laboratory for Genomic Medicine, Farmington, Connecticut, USA and stored at −80°C until DNA extraction or cultivation.

### mWGS DNA extraction

Metagenomic whole genome shotgun sequencing (mWGS) allows for comprehensively sample all genes in all organisms present in a given complex sample and to identify bacteria, viruses and fungi. Genomic DNA was extracted from frozen samples in the manual of the GenElute Bacterial DNA Isolation kit (Sigma Aldrich, NA2110-1KT, St. Louis, MO, USA) with the following minor modifications as previously described^60^: Briefly, 5 μl of Lysozyme (10 mg/ml, Sigma-Aldrich, L6876, St. Louis, MO, USA), 1 μl of Lysostaphin (5000 U/ml, Sigma-Aldrich, L9043, St. Louis, MO, USA), 1 μl mutanolysin (5000 U/ml, Sigma-Aldrich, M9901, St. Louis, MO, USA) were added to each sample for a digest at 37°C, 30 min. Samples were mechanically disrupted for 2x 3 min at 30 Hz (TissueLyser II, Qiagen, Hilden, Germany). This was followed by adding 5 μl of proteinase K (20 mg/ml, Sigma-Aldrich) and 300 μl of Solution C (55°C, 30 min). The samples were precipitated by adding 300 μl of 100% EtOH (Fisher Scientific, Fair Lawn, NJ, USA) and the lysates were loaded on the GenElute columns. Subsequent steps were carried out according to the manufacturer’s instructions.

### mWGS DNA sequencing

Libraries were prepared using the Nextera XT DNA Library Prep Kit (Illumina, San Diego, CA, USA) according to the manufacturer’s instructions with the difference of using one quarter of each reaction volume. Whole genome sequencing (WGS) was predominantly carried out using a 2x 150bp (paired end) sequencing protocol for the Illumina the NovaSeq 6000 Sequencing System (Illumina, San Diego, CA, USA) according to the manufacturer’s manual. Sequencing was conducted at the Genome Technologies core facility at the Jackson Laboratory for Genomic Medicine, Farmington, USA.

### Positive and negative controls

Per patient, one air sample (negative control) was collected and extracted as described above. Samples that yielded in a Qubit measurement were processed for library preparation and sequenced as described. Per extraction round, one sample of a defined, in-house mock community (25 diverse Gram-positive and Gram-negative bacteria and fungi) and a negative reagent control (nuclease-free water, Qiagen, Hilden, Germany) were included. For library generation, a negative control (nuclease-free water, Qiagen, Hilden, Germany) was included as well. One mock community sample was added to each sequencing run and a library/ extraction negative control was sequenced if a library product was measurable on the Qubit 2.0 Fluorometer (Thermo Fisher Scientific, Waltham, Massachusetts, USA) or identified on the 4200 TapeStation System (Agilent Technologies, Santa Clara, CA) using the High Sensitivity D1000 ScreenTape Assay (Agilent Technologies, Santa Clara, CA).

### mWGS data processing for taxonomic classification

The mWGS data analysis includes both taxonomic and functional profiling from complex microbial communities. After demultiplexing, the fastq files were processed with MetaPhlAn 4^61^ using the flags --add_viruses and --unclassified_estimation. Samples with no reads mapping to any taxa were excluded.

Relative proportions were used for all analyses. All taxonomic features at the species level with a mean relative abundance of 0.1% (denoise function^62^ across the dataset were removed from the dataset to reduce potential false positives and allow for multiple hypothesis correction (except for the comparison with the culturomics data).

### Isolate culturing and identification

Oral and nares samples from the patients destined for culturing were thoroughly vortexed, and then diluted 1:100 and 1:1000 in R2A, to increase the chance of recovering single colonies. 50µL from each dilution was then spread on half of an agar plate (R2A, LB, TSA blood agar, Chocolate agar, aerobic only: SaSelect) for each cultivation condition using a sterile spread tool (Thomas Scientific, 229616) as previously described^28^. Briefly, for agar cultivation, the following media were sourced from Fisher Scientific: Luria Broth (LB) agar (BP1425500), R2A agar (R454372), Tryptic Soy Broth (TSB) (DF0370-17-3), Chocolate agar (B21169X), Tryptic Soy Agar (TSA) with 5% sheep blood agar (B21261X). SaSelect agar plates (63748) were obtained from BioRad. The anaerobic atmosphere consisted of 5% hydrogen, 5% carbon dioxide, 90% nitrogen (Airgas, Z03NI9022000008). Aerobic cultures were conducted in ambient atmosphere.

Microbial isolates were identified as previously described^28^ using matrix assisted laser desorption ionization-time of flight (MALDI-TOF, Bruker Daltonics, Germany) mass spectrometry. Rapid DNA extraction from microbial isolates was adapted from Köser *et al*.^63^ and sequenced as previously described^28^ on the NovaSeq 6000 Sequencing System (Illumina, San Diego, CA, USA), 2×150bp paired end to approximately 100X coverage per genome. Following sequencing, genomic reads were preprocessed and dereplicated using PRINSEQ lite^64^ and trimmed using trimmomatic^65^. Reads were assembled into contigs using SPAdes^66^. QUAST^67^ was used to assess contig quality. Taxonomic classification was assigned to the contigs and the contigs were placed in a phylogenetic tree using GTDB^68,69^. Reference genomes were used for *Candida glabrata*, *S*. *hominis*, and *L*. *rhamnosus*. The phylogenetic tree was visualized using the R package ggtree^70^.

### Statistics and sample comparisons for metagenomics and culturomics data

Overlap of metagenomics and culturomics data was calculated extracting the uniquely identified species of genera in the culturomics data and metagenomics data. Data was analyzed and visualized using the following R packages: reshape2^71^, ggplot2^72^, tidyverse^73^, ggpubr^74^, dplyr^75^, plyr^76^, and ggvenn^77^.

### Comparative genomic analysis

Microbial genomes were annotated for virulence factors by blasting them against VFDB’s protein database A (verified virulence factors)^43^ using UBLAST^1^. A minimum e-value of 10^-5^ and percent identity of at least 80% was required. If multiple hits were identified, the hit with the lowest e-value was selected.

KEGG orthologs were annotated by blasting the microbial genomes against UniRef90 using UBLAST. A minimum e-value of 10^-9^ was required. If multiple hits were identified, the hit with the lowest e-value was selected. FMAP^44^ was then used to calculate the abundances (RPKM) of the KEGG orthologs with a minimum percent identity of 90%, identify differentially abundant genes between interferon non-stimulator and stimulator microbes, and determine KEGG pathways, modules, and operons that were enriched with the differentially abundant orthologs. Mann-Whitney U test was used to determine statistical significance (FDR adjusted P-value < 0.05).

Microbial gene names were annotated using eggnog-mapper^1^ and Prodigal^1^. HAllA^47^ was used to identify associations between microbial gene names and the host log_2_FC for each ISG for each microbial treatment. Associations were calculated using NMI (normalized mutual information) and the P-value was adjusted with FDR. Significance was set at an adjusted P-value < 0.05. The top 50 associations were visualized using HAllA’s built-in hallagram function.

### Air-liquid interface cell culture cultivation

9mm primary normal human bronchial epithelial air-liquid interface (ALI) tissue cultures were obtained from MatTek Corporation (EpiAirway AIR-100, Gothenburg, Sweden). The ALI cultures had been switched to antibiotic-free media 3 days prior to receipt to facilitate microbial growth. All batches were grown using cells from MatTek Corporation’s EpiAirway AIR-100 standard donor, a healthy adult male. ALI cultures were cultured according to the manufacturer’s directions. Briefly, upon arrival, the ALI cultures were placed in 6-well plates with 1mL of warmed antibiotic-free EpiAirway AIR-100 Maintenance Media (MatTek Corporation, Gothenburg, Sweden) or the equivalent EpiAirway AIR-100 Assay Media (MatTek Corporation, Gothenburg, Sweden) basally per well. The basal media was replaced daily, and the ALI cultures were kept at 37°C with 5% CO_2_.

### ALI treatments

For each microbial isolate, a single colony was grown overnight in sterile 1X Tryptic Soy Broth (TSB, Becton, Dickinson, and Company, #211825, Sparks, MD) with 0.1mg Vitamin K (Sigma Aldrich, #95271, St. Louis, MO) and 5mg heme / 1L TSB. For isolates that could not be grown from a single colony, a single colony was patched, and that patch was used to start a liquid culture. ODs were obtained by measuring absorbance at 600nM in a 96-well plate using Cytation Station 5 (BioTek Agilent Technologies, Santa Clara, CA). 10^8^ colony-forming units (CFUs) were taken from each liquid culture and washed with ultrapure water (Fisher Scientific, #AAJ71786AP, Hampton, NH) being resuspended in 100µL of ultrapure water (Fisher Scientific, #AAJ71786AP, Hampton, NH), with a final concentration of 10^7^ CFUs per 10µL.

Just prior to treatment, the ALI cultures were washed twice with transepithelial electrical resistance buffer (MatTek Corporation, Gothenburg, Sweden) to remove mucus. Each wash consisted of the addition and immediate removal of 400µL of the buffer. ALI cultures were then dosed with 10µL of microbial isolate/vehicle (ultrapure water, Fisher Scientific, #AAJ71786AP, Hampton, NH). Microbial isolates were dosed at 10^7^ CFUs. Dosed ALI cultures were incubated for 18 or 48 hours prior to harvest. Extra sample/vehicle were serially diluted in sterile PBS (MatTek Corporation, Gothenburg, Sweden) and grown on TSA with 5% sheep’s blood (Fisher Scientific, #221261, Hampton, NH) to determine the number of microbes added to the ALI cultures.

At harvest, 200µL of transepithelial electrical resistance buffer (MatTek Corporation, Gothenburg, Sweden) was added to the apical surface of each ALI culture, pipette mixed, and removed. 140µL of Buffer RLT (Qiagen, Hilden, Germany) + 1% beta-mercaptoethanol was added to each ALI culture, dissolving the tissue. The Buffer RLT-tissue solution was frozen at −80°C until RNA extraction. *S*. *epidermidis* Tü3298-GFP colonized ALI were visualized under blue light for a qualitative examination of colonization. The transepithelial electrical resistance buffer harvest wash was serially diluted in PBS (MatTek Corporation, Gothenburg, Sweden) and plated on TSA with 5% sheep’s blood (Fisher Scientific, #221261, Hampton, NH) or 1X TSB (Becton, Dickinson, and Company, #211825, Sparks, MD) with 1X Bacto Dehydrated Agar (Fisher Scientific, #214010, Hampton, NH) for CFU counts. Basal media was collected and frozen at −80°C for cytokine bead array assays.

### RNA extraction and RNA-seq

All RNA extraction and sequencing library preparation steps were performed in a sterile tissue culture hood. RNA was extracted using RNeasy 96 QIAcube HT kit (Qiagen, Hilden, Germany) according to the manufacturer’s directions. Samples were eluted in nuclease-free water (Qiagen, Hilden, Germany) and frozen at −80°C until sequencing preparation. RNA quality was evaluated using the 4200 TapeStation System (Agilent Technologies, Santa Clara, CA) with the High Sensitivity RNA ScreenTape Analysis (Agilent Technologies, Santa Clara, CA) or RNA ScreenTape Analysis (Agilent Technologies, Santa Clara, CA). RINs ranged from 1.7-9.9 with an average of 8.4 and a median of 8.9. RNA quantity was measured on the Qubit 2.0 Fluorometer (Thermo Fisher Scientific, Waltham, Massachusetts, USA). The sequencing libraries were prepared with NEBNext rRNA Depletion Kit v2 (New England Biolabs, Ipswich, MA) and NEBNext Ultra II Directional RNA Library Prep Kit for Illumina (New England Biolabs, Ipswich, MA) following the manufacturer’s directions. Library quality was evaluated using the 4200 TapeStation System (Agilent Technologies, Santa Clara, CA) with the High Sensitivity D1000 ScreenTape Assay (Agilent Technologies, Santa Clara, CA). Library quantity was measured on the Qubit 2.0 Fluorometer (Thermo Fisher Scientific, Waltham, Massachusetts, USA). Samples were sequenced using Illumina NovaSeq targeting 30 million reads. Reads per sample ranged from 76 thousand to 318 million with an average and median of 29 million reads.

### Transcriptional profiling, gene set enrichment analysis, and CFU confounder analyses

Raw RNA-seq reads was processed with trimmomatic 0.39^65^ to remove low-quality reads. QC’d reads were mapped to CHM13v2.0 reference genome using STAR 2.5.3a^78^. The raw count of reads aligned to each gene was computed using featureCounts (from Subread1.6.4)^79^. Differentially expressed genes were identified and expression level changes of each gene was computed using DESeq2^80^. Gene set enrichment analysis was performed on pre-ranked gene lists calculated from output of DESeq2 on ALI transcriptomes by GSEA4.3.2 or clusterProfiler^81,82^. CFU confounder analyses were generated by MaAsLin2^32^ with TSS (total sum-scaling) normalization. Fixed effects included CFUs, ALI batch, and timepoint. An effect was considered a confounder if the FDR adjusted P-value < 0.05.

ALI transcriptomes were analyzed with CFU counts of microbial treatments. Genes were considered as confounded by CFU if counts of genes are significant associated with CFU counts within the dataset.

### Gene list generation and determination of responsive genes in RNA-seq data

ISG gene lists were adapted from the literature: total ISG list^83^, 6-gene ISG list^34^, and 30-gene ISG list^35^. The antibacterial gene list was derived from MSigDB^36^ with gene ontologies^37,38^ (GO:0005125, GO: 0061844, GO: 0019730, GO: 0050832, GO: 0009620). The keratin gene list was adapted from the literature^41^. The junction and mucin gene lists was derived from HGNC (HUGO Gene Nomenclature Committee)^42^. The junction gene list was further adapted from the literature^40^. To generate responsive genes within gene lists above, genes of low transcripts were filtered out. Within each gene list, genes were further processed by removing genes with raw read count lower than 5. The remaining genes were further processed by hierarchical clustering which implanted method in seaborn Python package^84^. Clusters of genes that were enriched with increased expression, as determined by high log_2_FC, were defined as responsive genes. The degree of increased expression depended on the gene list.

### Statistics and sample comparisons for RNA-seq data

Data was analyzed and visualized using the following Python packages: seaborn^84^, scikit-learn^85^, statannot^86^, pandas^87^, numpy^88^, and matplotlib^89^; and R packages: ComplexHeatmap^90^, reshape2^71^, and ggplot2^72^.

### Conditioned supernatant for cytokine induction/repression by Cytometric Bead Array (CBA)

Each cultured microbe was grown overnight in TSB with 0.1mg vitamin K and 5mg heme / 1L TSB in a 96-well deep well plates (#503501, Southern Labware, Cumming, GA, USA). ODs were obtained by measuring absorbance at 600nM in a 96-well plate using Cytation Station 5 (BioTek Agilent Technologies, Santa Clara, CA). The liquid cultures were centrifuged at 3500 rpm for 5 minutes (Sorvall Legend X1R M20 rotor, Thermo Scientific, Waltham, MA, USA) to separate the bacterial pellet and bacterial supernatant. The bacterial supernatant was collected and sterile filtered (0.2μm) using a filter plate (#MSGVS2210, Sigma Aldrich, St. Louis, MO, USA) at 3500 rpm for 5 minutes (Sorvall Legend X1R M20 rotor, Thermo Scientific, Waltham, MA, USA). The sterile bacterial supernatants were stored at −80°C until subsequent use.

Lung epithelial cell line A549 were stimulated overnight with sterile (0.2μm filtered) bacterial supernatants. Briefly, 1×10^5^ cells per well were with 20% volume/volume of each supernatant overnight. The conditioned media was collected for CBA analysis of human inflammatory cytokines concentration (IL-8, IL-6, IL-1, IL-12, IL-10, and TNF). Six bead populations with distinct fluorescence intensities were coated with capture antibodies specific for IL-8, IL-6, IL-10, IL-1, TNF, and IL-12p70 proteins (Becton, Dickinson, and Company BioSciences, #551811, Sparks, MD). They were incubated together with the samples (sterile bacterial supernatant or microbially colonized ALIs’ basal media) or recombinant standards, and PE-conjugated detection antibodies, to form sandwich complexes and were detected by flow cytometry. Results were generated in graphical and tabular format using the BD CBA Analysis Software (Becton, Dickinson, and Company BioSciences, Sparks, MD). The standard curve for each protein covers a defined set of concentrations from 20 – 5000 pg/ml.

Data was analyzed and visualized using the following R packages: ggplot2^72^, ggpubr^74^, dplyr^75^, rstatix^91^, tidyverse^73^, forcats^92^, and RColorBrewer^93^.

### Immunofluorescence staining

ALI cultures were washed with 1X PBS (MatTek Corporation, Gothenburg, Sweden), then embedded in OCT (Sakura Finetek USA, Torrance, CA, USA) and snap-frozen at −80°C. Frozen sections were cut at 8 µm, air dried on Superfrost plus slides, fixed again with 4% paraformaldehyde (#28906, Thermo Fisher Scientific, Waltham, MA, USA) for 15 minutes and permeabilized with 1X PBS/0.1% Triton-X (#HFH10, Thermo Fisher Scientific, Waltham, MA, USA) for 15 min. Tissue sections were treated with Fc Receptor Block (#NB309, Innovex Bioscience, Richmond, CA, USA), followed by Background Buster (#NB306, Innovex Bioscience, Richmond, CA, USA). The sections were then stained with anti-MX1 primary antibody (polyclonal Rabbit N2C2, #GTX110256, GeneTex, Irvine, CA, USA) for one hour followed by secondary antibody (anti-rabbit IgG AF555, #406412, BioLegend, San Diego, CA, USA) for 30 min at room temperature in 1X PBS/5% BSA/0.05% saponin. Then, sections were washed three times with 1X PBS for 15 min. Respective isotype antibodies were used as controls. Finally, sections were counterstained with 1 µg/ml of 4’,6-diamidino-2-phenylindole (DAPI, #D1306, Thermo Fisher Scientific, Waltham, MA, USA) then, mounted with Fluoromount-G (#00-4958-02, Thermo Fisher Scientific, Waltham, MA, USA), acquired using a Leica SP8 confocal microscope (Leica Microsystems, Wetzlar, Germany) for high resolution images and Thunder widefield microscope (Leica Microsystems, Wetzlar, Germany) for histocytometry both with Leica LAS X software (Leica Microsystems, Wetzlar, Germany) and analyzed using Imaris software (Bitplane, Oxford Instruments, Abingdon, United Kindgom).

Data was analyzed and visualized using the following R packages: ggplot2^72^, ggpubr^74^, dplyr^75^, rstatix^91^, and RColorBrewer^93^. All results are expressed as a mean of the replicates, unless specified. All comparisons were made between infection conditions with time point-matched, uninfected controls.

### Transepithelial electrical resistance (TEER) measurements

TEER measurements were taken using EVOM Manual for TEER Measurement (#EVM-MT-03-01, WPI, Sarasota, FL, USA) and STX4 EVOM Electrode (#EVM-EL-03-03-01, WPI, Sarasota, FL, USA) following 18 and 48 hours of colonization. The electrodes were equilibrated following manufacturer’s instructions. ALIs were transferred to a 12-well plate. 1mL of media was added to the basal side and 300uL of TEER buffer was added to the apical surface. Three readings were taken of each insert, even spread across the insert. In between each reading, the electrodes were washed in 70% ethanol and then washed in TEER buffer.

Data was analyzed and visualized using the following R packages: ggplot2^72^, ggpubr^74^, dplyr^75^, rstatix^91^, and RColorBrewer^93^.

---

**Figure S1.**
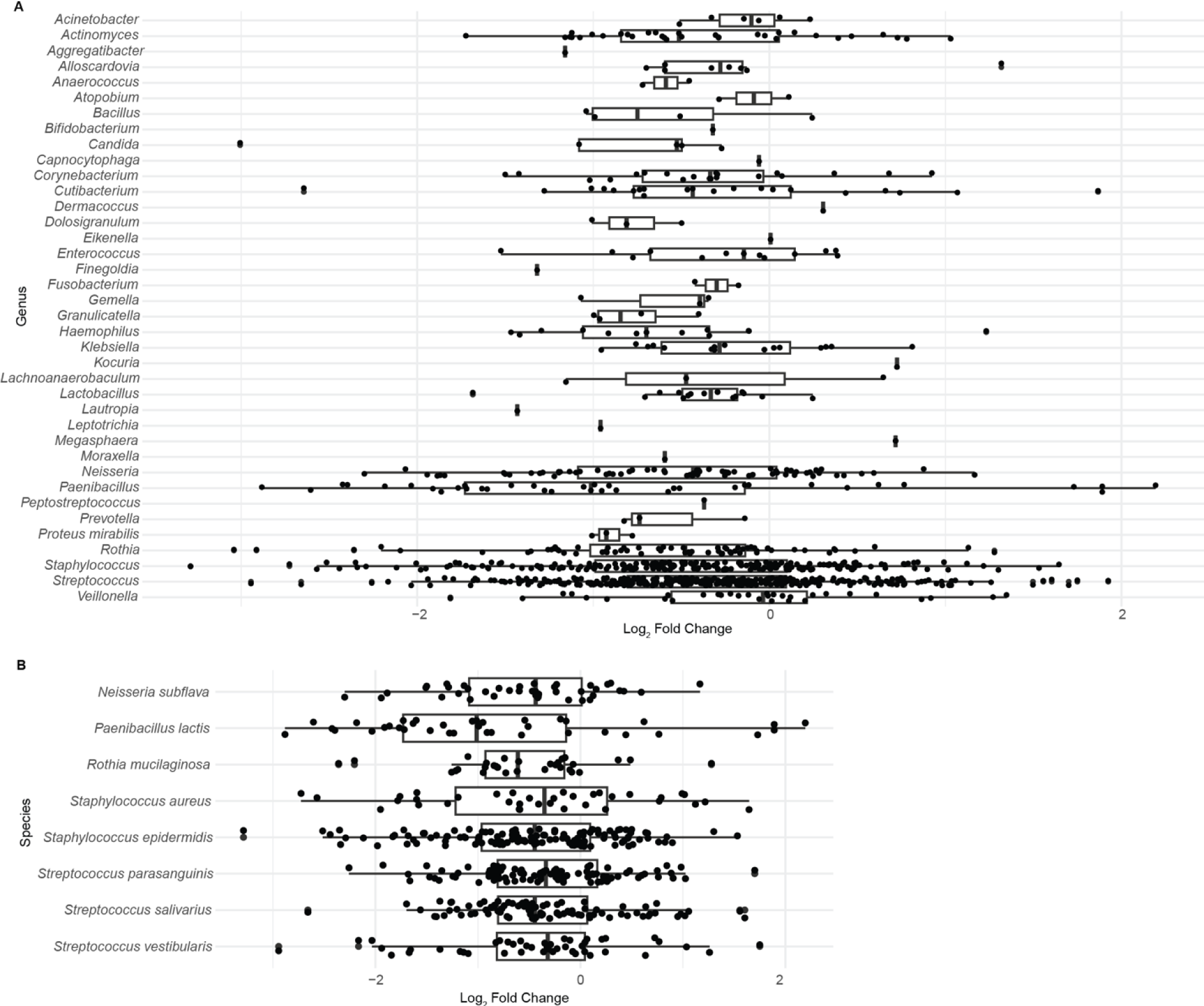
Secreted microbial metabolites modulate IL-8 secretion independent of phylogeny in 2D lung cultures. Microbial supernatant from isolated microbes was applied to A549 epithelial adenocarcinoma cells and secreted IL-8 (as well as other cytokines, which showed no induction/repression) was quantified using cytokine bead assay. **A)** Log_2_ fold change (FC) in secreted IL-8 (relative to vehicle treated cells) for each microbe, excluding those without an ID, organized by genus. **B)** Log_2_FC in secreted IL-8, organized by species, for species with at least 30 strains.

**Figure S2.**
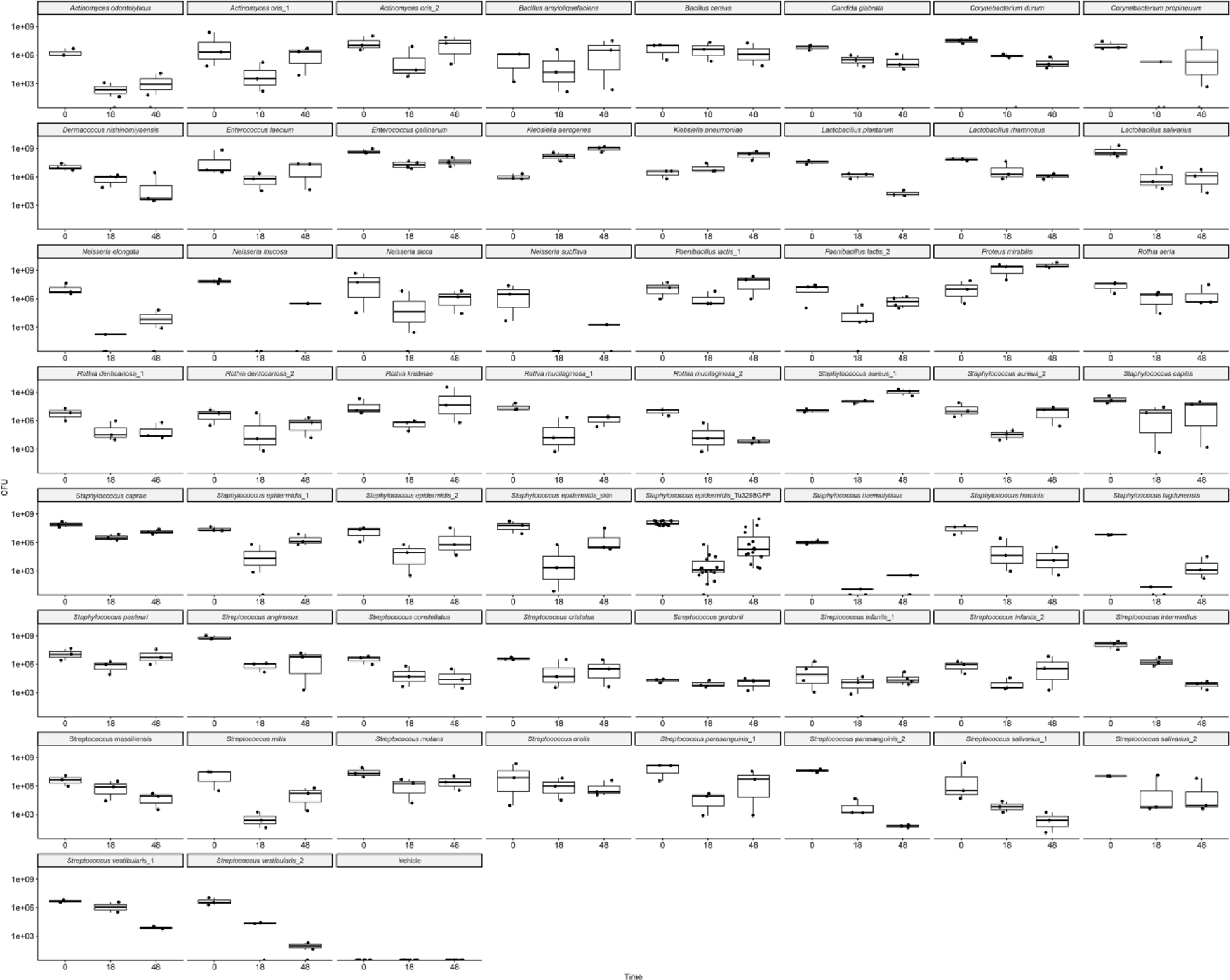
Microbial CFUs at dosing, 18 hours, and 48 hours. At each harvest, the colonizing microbes/vehicle were washed off of the ALIs and, along with the dosing microbes/vehicle, serially diluted and grown on tryptic soy agar (TSA) or TSA with 5% sheep’s blood plates to quantify colony forming units (CFUs).

**Figure S3.**
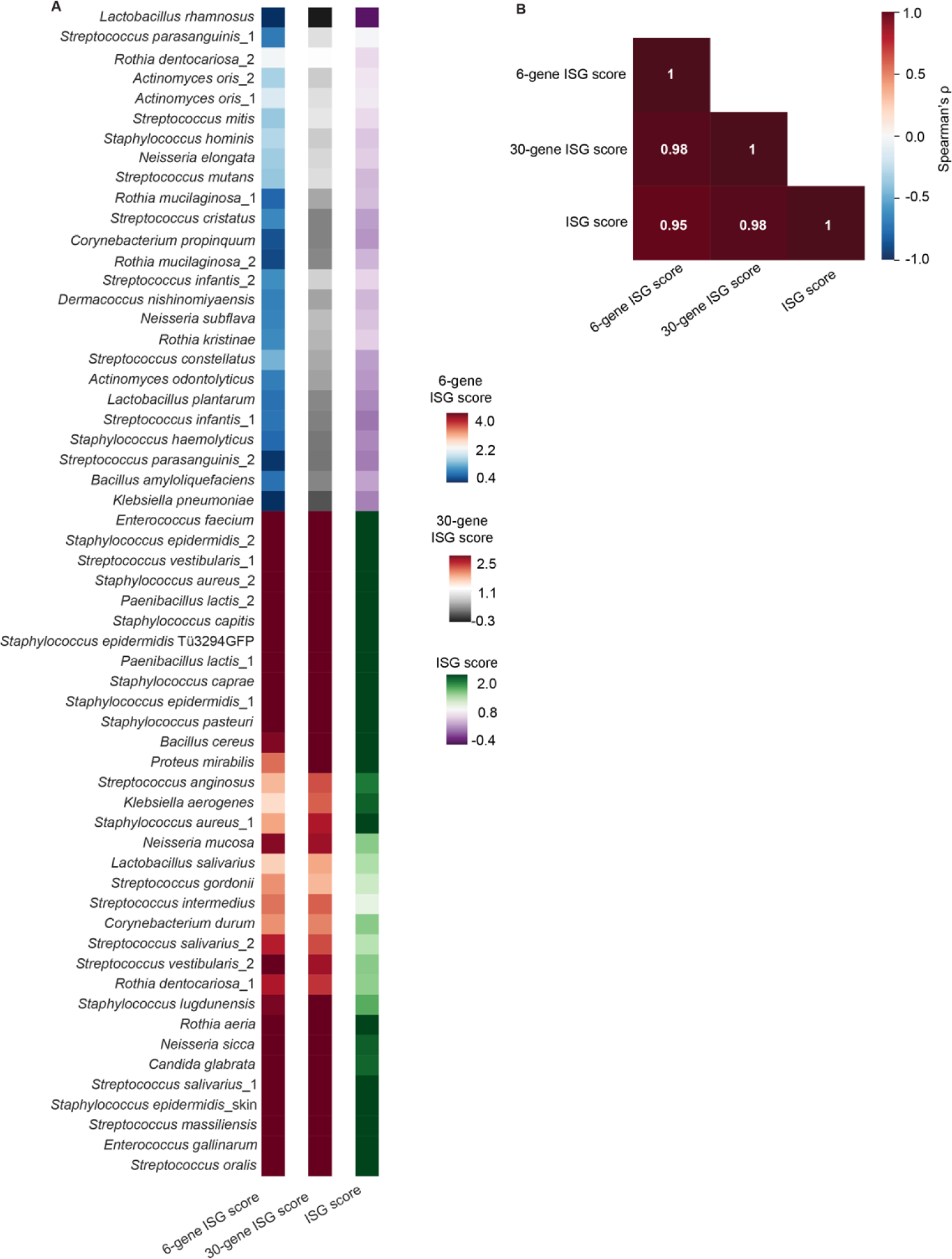
Comparison of ISG score used in this analysis with published 6-gene and 30-gene ISG scores. **A**) Heatmaps comparing each microbe’s ISG score when calculated with our ISG panel, versus previously published 6-gene and 30-gene ISG scores. **B**) Correlation (Spearman’s coefficient) between the three ISG scores.

**Figure S4.**
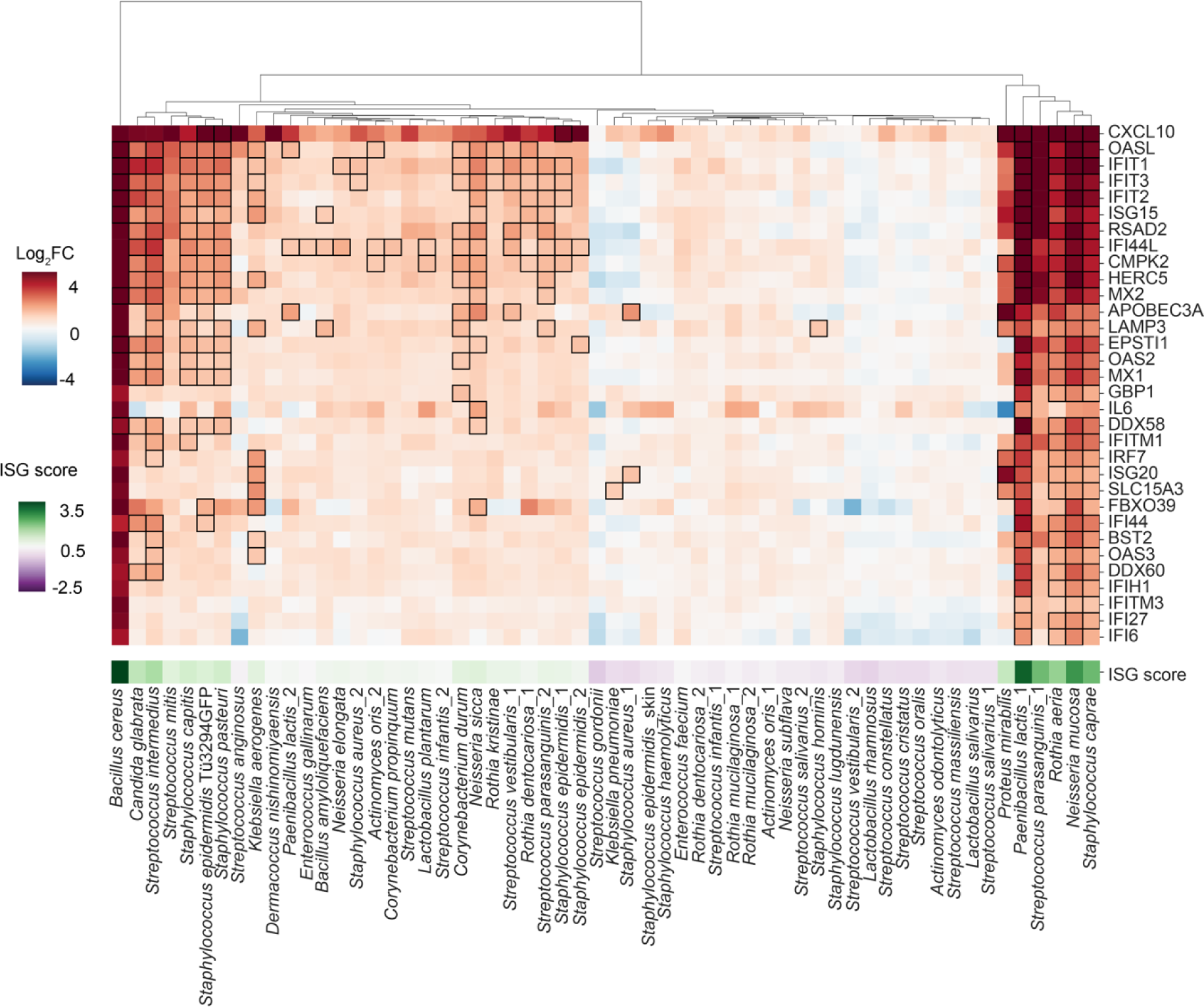
ISG gene expression changes following 18 hours of colonization. Heatmap (log_2_FC) of ISGs following 18 hours of colonization, relative to vehicle. Microbial treatment is hierarchically clustered. Black boxes indicate statistical significance (FDR adjusted P-value < 0.1). Below the heatmap is the microbe’s ISG score based on the 18 hour heatmap.

**Figure S5.**
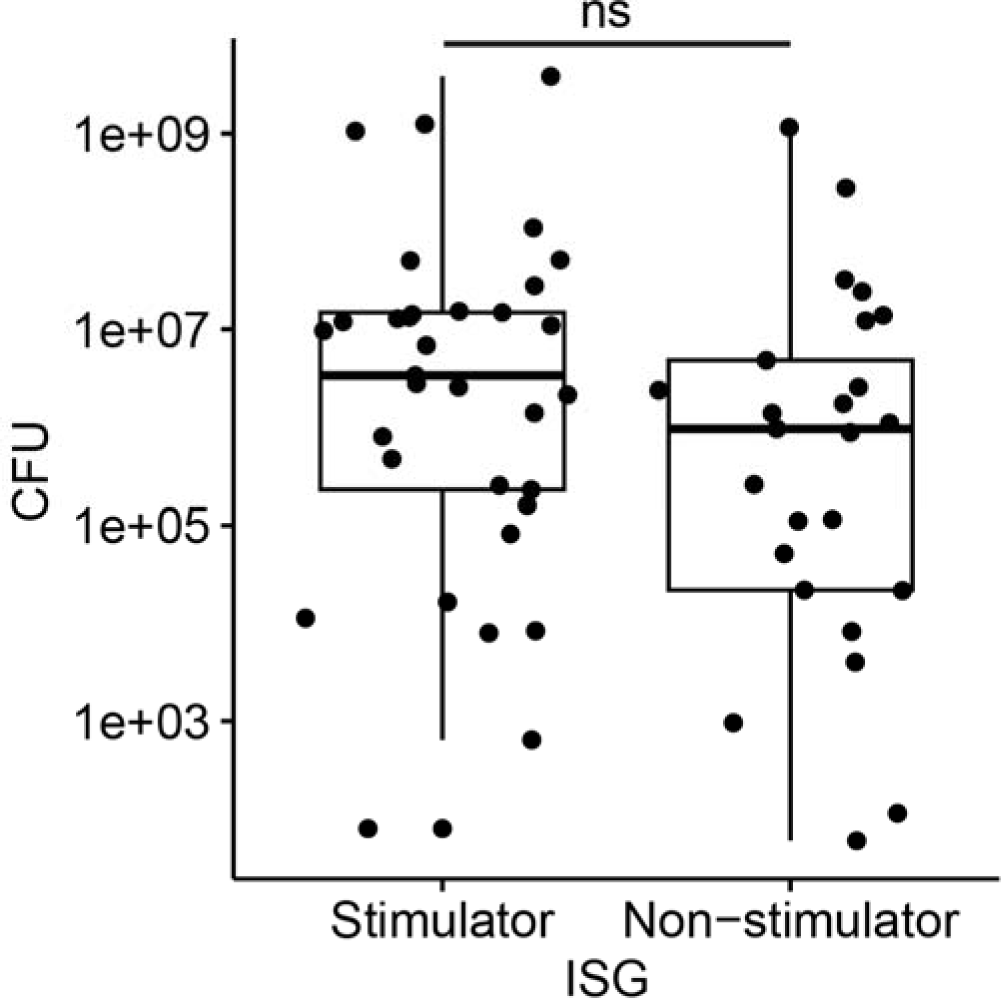
ISG stimulator versus non-stimulator ability is not dependent upon microbial load. Boxplot of each microbial’s CFUs at 48 hours grouped by the microbe’s ISG stimulator versus non-stimulator category; box edges indicate the 25^th^ and 75% percentiles. Each point is the average of the three replicates to avoid pseudoreplication. Statistical analysis is two-sided Mann-Whitney U test. ns indicates not significant.

**Figure S6.**
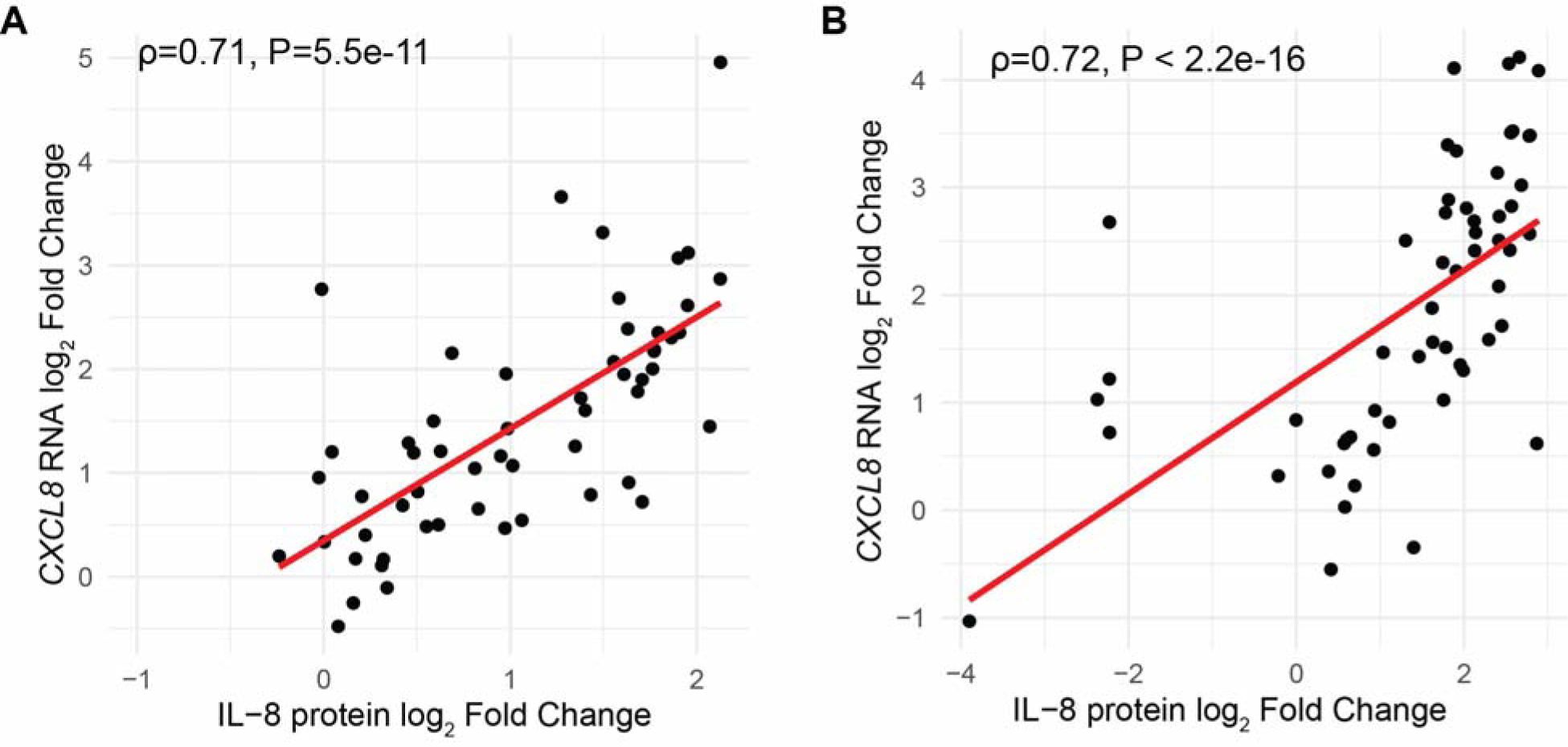
Basal secretion of IL-8 correlates with *CXCL8* gene expression changes at 18 and 48 hour timepoints. Correlation plots (Spearman’s correlation coefficient) of log_2_FC of *CXCL8* (IL-8) gene expression relative to vehicle (y-axis) and average (n=3) basally secreted IL-8 log_2_FC relative to vehicle (x-axis) at **A**) 18 hours and **B**) 48 hours.

**Figure S7.**
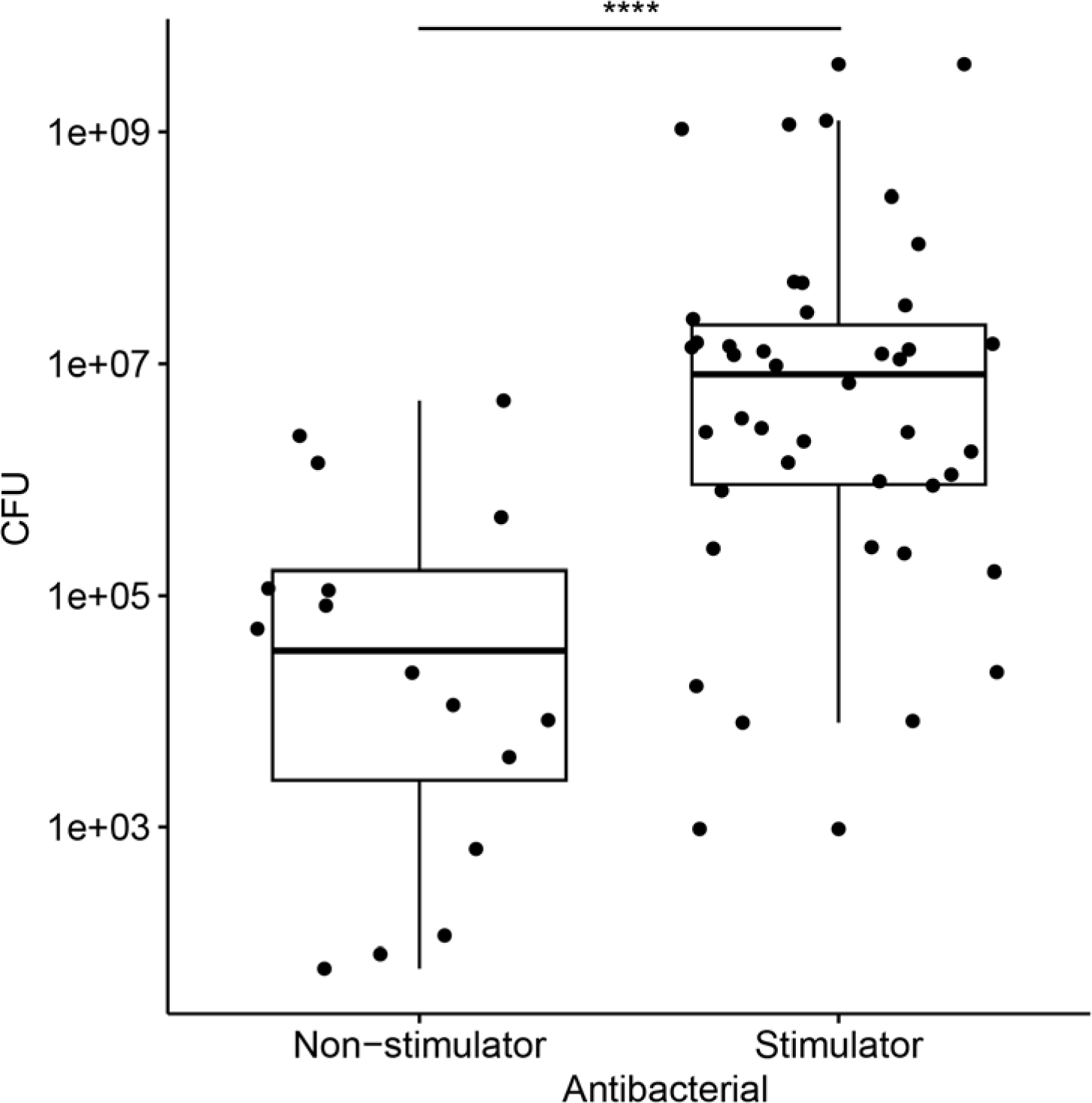
CFUs in part drive microbial antibacterial non-stimulator versus stimulator category. Boxplot of each microbial treatment’s CFUs (average of n=3) at 48 hours separated by antibacterial non-stimulator versus stimulator category; box edges indicate 25^th^ and 75^th^ percentiles. Statistical analysis is two-sided Mann-Whitney U test with Bonferroni correction. * indicates P-value <0.1, **<0.05, ***<0.01, ****<0.001.

**Figure S8.**
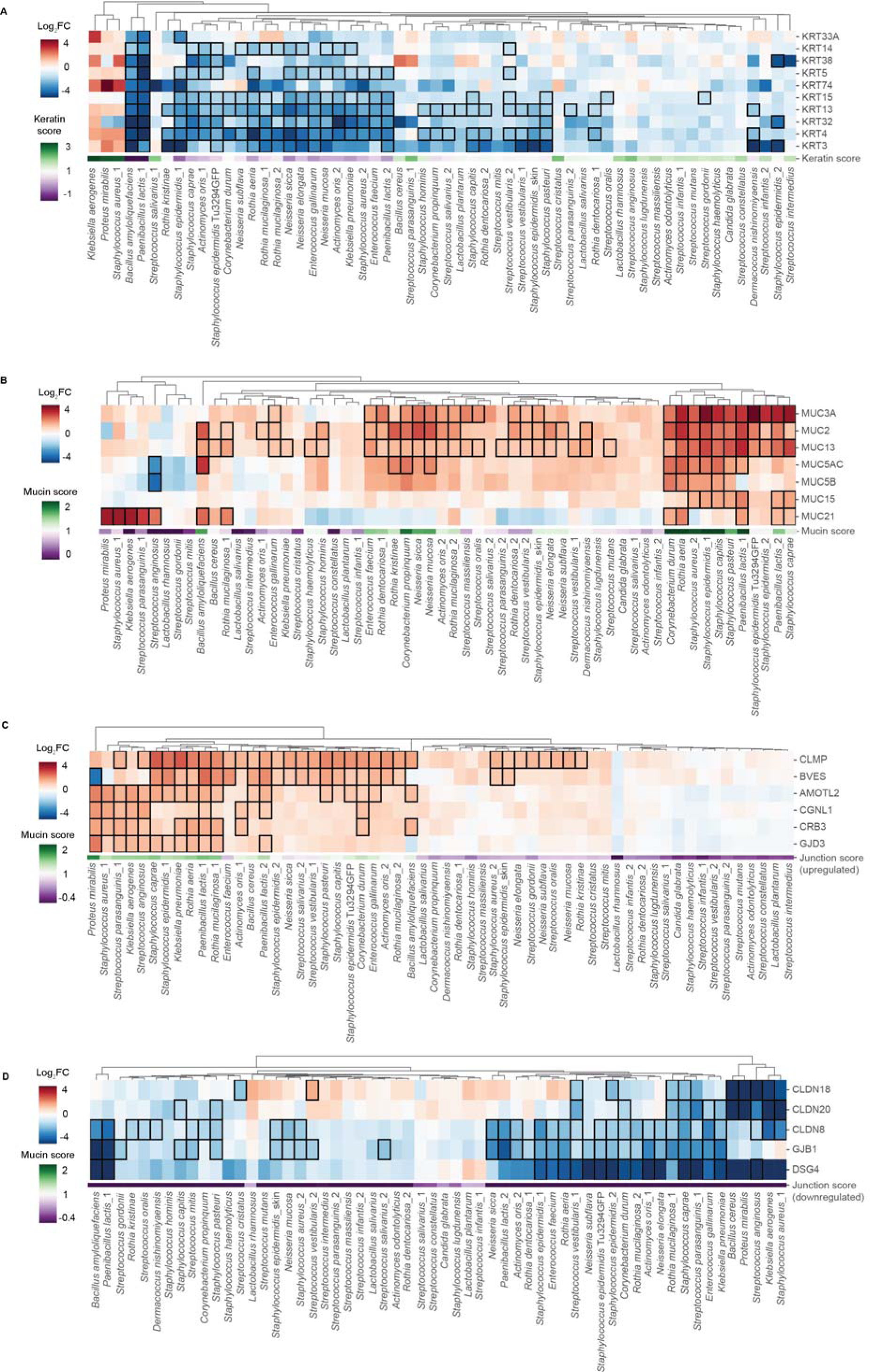
Keratin and mucin gene list gene expression change at 48 hour timepoint. Heatmap (log_2_FC) of **A**) keratin, **B**) mucin, and **C**) junction (tight, adherens, and gap) genes following 48 hours of microbial colonization relative to vehicle. Black boxes indicate statistical significance (FDR adjusted P-value < 0.1). Microbial treatments are hierarchically clustered. Below the heatmap is the respective score.

